# IDSL.CCDB: a database for exploring inter-chemical correlations in metabolomics and exposomics datasets

**DOI:** 10.1101/2022.02.01.478739

**Authors:** Dinesh Kumar Barupal, Priyanka Mahajan, Sadjad Fakouri Baygi, Robert O Wright, Manish Arora, Susan L. Teitelbaum

## Abstract

Inter-chemical correlations in metabolomics and exposomics datasets provide valuable information for studying relationships among reported chemicals measured in human specimens. With an increase in the size of these datasets, a network graph analysis and visualization of the correlation structure is difficult to interpret. While co-regulatory genes databases have been developed, a similar database for metabolites and chemicals have not been developed yet. We have developed the Integrated Data Science Laboratory for Metabolomics and Exposomics - Chemical Correlation Database (IDSL.CCDB), as a systematic catalogue of inter-chemical correlation in publicly available metabolomics and exposomics studies. The database has been provided via an online interface to create single compound-centric views that are clear, readable and meaningful. We have demonstrated various applications of the database to explore: 1) the chemicals from a chemical class such as Per- and Polyfluoroalkyl Substances (PFAS), polycyclic aromatic hydrocarbons (PAHs), polychlorinated biphenyls (PCBs), phthalates and tobacco smoke related metabolites; 2) xenobiotic metabolites such as caffeine and acetaminophen; 3) endogenous metabolites (acyl-carnitines); and 4) unannotated peaks for PFAS. The database has a rich collection of 36 human studies, including the National Health and Nutrition Examination Survey (NHANES) and high-quality untargeted metabolomics datasets. IDSL.CCDB is supported by a simple, interactive and user-friendly web-interface to retrieve and visualize the inter-chemical correlation data. The IDSL.CCDB has the potential to be a key computational resource in metabolomics and exposomics facilitating the expansion of our understanding about biological and chemical relationships among metabolites and chemical exposures in the human body. The database is available at www.ccdb.idsl.me site.

## Introduction

Combined exposures to millions of different chemicals and its impact on the health and development of human body is a major component of the exposome.^1^ The chemical exposome is made up of nutrients and environmental non-food chemicals, consisting of natural and synthetic exogenous compounds.^2–4^ After entering the body through biotransformation they form the metabolome, which includes metabolic end products of the host and its commensal microbiota. This chemical space (e.g. industrial chemicals, nutrients, drugs, and bioactive internal molecules such as hormones and oxylipins) has significant influence on health trajectories and chronic health outcomes and is implicated in all diseases, including cancer as well as neurological, cardiovascular, and respiratory diseases^5–15^. Emerging evidence demonstrates that the scale, magnitude, and structural diversity^3, 16^ of the internal chemical space is vast and that many chemicals could be classified together because they are structurally and functionally related to each other^17–19^. A systematic understanding and cataloging of targeted and untargeted analyses of small molecules measured in biospecimens is needed, as such datasets are critical to translate the information gathered from exposomics and metabolomics projects^20^. These key datasets include: 1) population-scale biomonitoring surveys; 2) targeted analysis of multiple analytes in hypothesis-driven studies (typically 10-100); and 3) untargeted analysis of thousands of chemicals using a high-resolution mass spectrometry instrument^21, 22^. They cover key high priority exposome chemicals^23^ including carcinogens^24, 25^, endocrine disrupters^26^ and industry chemicals^27^. These core datasets support different statistical and bioinformatics analyses to reveal novel risk factors, hidden metabolic pathways, detrimental exposures and biomarkers for disease.

Computing the correlation coefficient between intensities of two chemicals is a fundamental statistical approach classically used to study enzyme kinetics^28^ and biotransformation^29^. For modern multi-analyte targeted and untargeted assays, a pair-wise correlation matrix among detected chemicals is computed for almost every study because this matrix can be used to assess chemical clustering^30^, peak annotation^31^, heatmaps^32^, and correlation network visualization^30^. Correlation among gene expression data is often interpreted as evidence of a co-regulatory pathway such as a common transcription factor that controls expression of a group of genes^33, 34^. As a corollary, with chemicals correlation can reflect common exposure origins^35^ as well as shared metabolic regulations, such as absorption pathways, biotransformation^36, 37^ and elimination as seen in drugs and their metabolic products^38, 39^. For exposomic projects, the probable interpretation of inter-chemical correlations is summarized in Figure 1. The biological interpretation covers both kinetics (i.e. the metabolic fate of a chemical^40^) and dynamics (i.e. the toxic effect of chemical exposure). The system connects to key metabolic pathways^41^, and creates logical groupings of similar exposures in a chemical class^30^. It can also indicate that two chemicals share an exposure source, such as occupation, consumer products^42^, or food^43^. Despite the utility and application of inter-chemical correlation data, a database of these inter-chemical correlations has not yet been developed. Our methods therefore fill several research gaps^44, 45^ and will inform chemical regulation policies while expanding understanding of fundamental metabolic processes^46^ that transform exposome chemicals.

**Figure 1.**
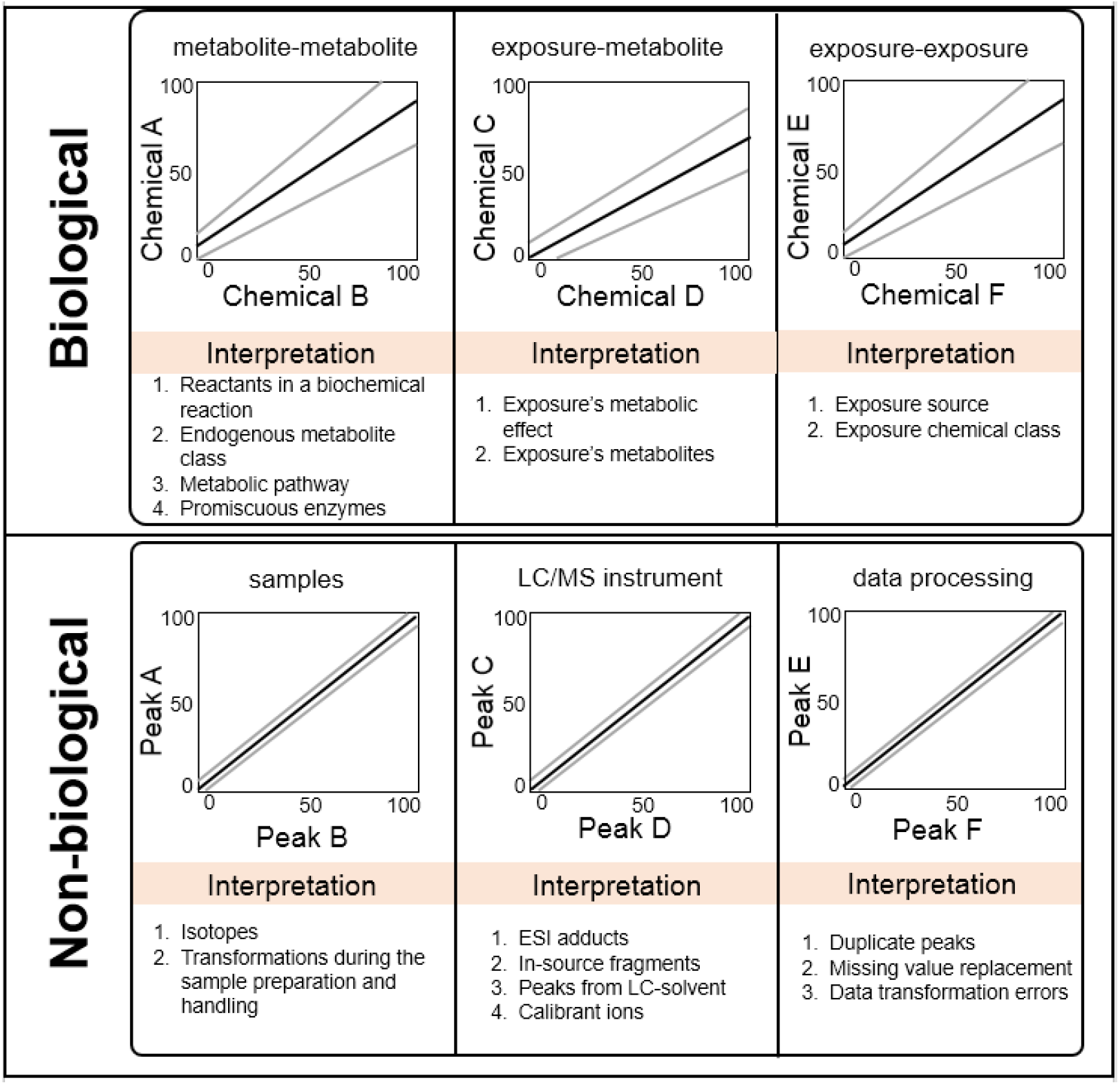
Probable interpretation of correlation in targeted and untargeted LC/HRMS datasets.

Metabolomic correlation network analyses show that chemically similar compounds and compounds belonging to the same pathway tend to show a higher clustering coefficient^47–49^. However, creating and analyzing those networks for large and comprehensive metabolomics datasets that often have over ten thousand reported peaks is computationally challenging. It is even more difficult to create and analyze such network graphs for metabolomics datasets that are generated using multiple LC/GC assays (e.g. RP(+) and HILIC(−) modes) for hundreds of samples^50^. There is a need to catalogue these correlations in a systematic database for mining them in various interpretational contexts.

Herein, we describe a new database, IDSL.CCDB, which catalogues pair-wise inter-chemical correlations from publicly available metabolomics and exposomics studies. It is the largest database of pairwise correlations to date and provides new opportunities for interpreting metabolomics datasets for structural and biological relationships. The database is publicly available at www.ccdb.idsl.me.

## Methods

### Selection of studies

Table 1 provides the list of studies and the details about the number of compounds and samples. For the development of the database, we constrained our approach to human specimen studies having at least 50 samples. To include a study in the IDSL.CCDB, the data were re-formatted into IDSL.CCDB Excel template (SI File 1). The template requires three sheets 1) “data_dictionary” which contains the metadata for annotated and unannotated compounds 2) “data_matrix” which contains the intensity data for all peaks and 3) “sample_metadata” which contains the information about each sample. If data from different chromatography and ionization modes were available, data were stacked in the “data_dictionary” and “data_matrix” sheets. If data were not scaled or normalized, we applied a log2 transformation before computing the correlation.

**Table 1:** Covered studies in the IDSL.CCDB on January 1^st^ 2022.

| S.No | Accession ID | Title | Number of Samples | Number of Peaks | Specimen |
| --- | --- | --- | --- | --- | --- |
| 1 | ST000923 | Longitudinal Metabolomics of the Human Microbiome in Inflammatory Bowel Disease<sup>51</sup> | 546 | 81867 | Stool |
| 2 | ST001000 | Gut microbiome structure and metabolic activity in inflammatory bowel disease<sup>52</sup> | 220 | 8847 | Stool |
| 3 | ST001192 | A library of human gut bacterial isolates paired with longitudinal multiomics data enables mechanistic microbiome research<sup>53</sup> | 180 | 54402 | Stool |
| 4 | ST001520 | Stool unknowns profiled using hybrid nontargeted methods (part-II)<sup>54</sup> | 166 | 54014 | Stool |
| 5 | NHANES | National Health and Nutrition Examination Survey – USA. Continuous NHANES data from 1999-2020 period.<br>https://www.cdc.gov/nchs/nhanes/index.htm | 107,258 (SEQN IDs) | 1,721 (variables) | Blood/Urine |
| 6 | ST001223 | Host Metabolic Response in Early Lyme Disease<sup>55</sup> | 518 | 2193 | Blood |
| 7 | ST001081 | Combined NMR and MS Analysis of PC patient serum (part-I)<sup>56</sup> | 168 | 459 | Blood |
| 8 | ST001082 | Combined NMR and MS Analysis of PC patient serum (part-II)<sup>56</sup> | 265 | 24928 | Blood |
| 9 | MTBLS1684 | A multi-omic analysis of birthweight in newborn cord blood reveals new underlying mechanisms related to cholesterol metabolism<sup>57</sup> | 499 | 63393 | Blood |
| 10 | ST001682 | Untargeted urine LC-HRMS metabolomics profiling for bladder cancer binary outcome classification | 311 | 982 | Urine |
| 11 | MTBLS136 | Serum metabolomic profiles associated with postmenopausal hormone use<sup>58</sup> | 1336 | 1385 | Blood |
| 12 | MTBLS204 | Metabolomics analysis of human acute graft-versus-host disease reveals changes in host and microbiota-derived metabolites<sup>59</sup> | 86 | 801 | Blood |
| 13 | MTBLS205 | Metabolomics analysis of human acute graft-versus-host disease reveals changes in host and microbiota-derived metabolites<sup>59</sup> | 112 | 929 | Blood |
| 14 | ST001516 | Identification of distinct metabolic perturbations and associated immunomodulatory events during intra-erythrocytic development stage of pediatric Plasmodium falciparum malaria<sup>60</sup> | 199 | 668 | Blood |
| 15 | ST001517 | Identification of distinct metabolic perturbations and associated immunomodulatory events during intra-erythrocytic development stage of pediatric Plasmodium falciparum malaria<sup>60</sup> | 106 | 652 | Blood |
| 16 | ST001639 | Plasma Metabolomic signatures of COPD in a SPIROMICS cohort<sup>61</sup> | 649 | 1174 | Blood |
| 17 | ST001212 | Fish-oil supplementation in pregnancy, child metabolomics and asthma risk<sup>62</sup> | 577 | 656 | Blood |
| 18 | ST001827 | The pregnancy metabolome from a multi-ethnic pregnancy cohort<sup>63</sup> | 410 | 1110 | Blood |
| 19 | IDSLCCDB00001 | Plasma and Fecal Metabolite Profiles in Autism Spectrum Disorder<sup>15</sup> | 222 | 1611 | Blood |
| 20 | IDSLCCDB00002 | Potential role of indolelactate and butyrate in multiple sclerosis revealed by integrated microbiome-metabolome analysis<sup>64</sup> | 180 | 517 | Blood |
| 21 | IDSLCCDB00003 | Comprehensive Circulatory Metabolomics in ME/CFS Reveals Disrupted Metabolism of Acyl Lipids and Steroids<sup>65</sup> | 52 | 1790 | Blood |
| 22 | IDSLCCDB00004 | Plasma Metabolomic Profiling in Patients with Rheumatoid Arthritis Identifies Biochemical Features Indicative of Quantitative Disease Activity<sup>66</sup> | 128 | 686 | Blood |
| 23 | IDSLCCDB00005 | Alterations in Polyamine Metabolism in Patients With Lymphangioleiomyomatosis and Tuberous Sclerosis Complex 2-Deficient Cells<sup>67</sup> | 78 | 1989 | Blood |
| 24 | IDSLCCDB00006 | Metabolic perturbation associated with COVID-19 disease severity and SARS-CoV-2 replication<sup>68</sup> | 72 | 1086 | Blood |
| 25 | IDSLCCDB00007 | Multi-omics of host-microbiome interactions in short- and long-term Myalgic Encephalomyelitis/Chronic Fatigue Syndrome (ME/CFS) | 184 | 1278 | Blood |
| 26 | ST001411 | Plasma metabolites of lipid metabolism associate with diabetic polyneuropathy in a cohort with screen-tested type 2 diabetes: ADDITION-Denmark<sup>69</sup> | 106 | 991 | Blood |
| 27 | ST001412 | Metabolomics study in Plasma of Obese Patients with Neuropathy Identifies Potential Metabolomics Signatures | 131 | 842 | Blood |
| 28 | ST001515 | A Metabolomic Signature of Glucagon Action in Healthy Individuals with Overweight/Obesity Humans<sup>70</sup> | 187 | 649 | Blood |
| 29 | ST001171 | Metabolomics of World Trade Center Exposed New York City Firefighters | 248 | 2504 | Blood |
| 30 | ST001430 | Metabolic dynamics and prediction of gestational age and time to delivery in pregnant women<sup>48</sup> | 781 | 9651 | Blood |
| 31 | ST001705 | Machine learning-enabled renal cell carcinoma status prediction using multi-platform urine-based metabolomics (part-I)<sup>71</sup> | 256 | 7097 | Urine |
| 32 | ST000292 | LC-MS Based Approaches to Investigate Metabolomic Differences in the Plasma of Young Women after Drinking Cranberry Juice or Apple Juice<sup>72</sup> | 51 | 3395 | Blood |
| 33 | ST000919 | Investigating Eicosanoids Implications on the Blood Pressure Response to Thiazide Diuretics | 140 | 10322 | Blood |
| 34 | ST000954 | Explore Metabolites and Pathways Associated Increased Airway Hyperresponsiveness in Asthma | 55 | 7930 | Blood |
| 35 | ST001231 | Plasma untargeted metabolomics study of pulmonary tuberculosis<sup>73</sup> | 159 | 17146 | Blood |
| 36 | MTBLS2295 | High-Precision Automated Workflow for Urinary Untargeted Metabolomic Epidemiology<sup>74</sup> | 87 | 655 | Urine |

### Processing of untargeted metabolomics studies

Only untargeted liquid chromatography high resolution mass spectrometry studies were selected. For each selected untargeted study (Table 1), we searched for a set of data types in the EBI-MetaboLights and MetabolomicsWorkBench repositories. The set included 1) raw data files in the vendor specific or open format such as mzML or mzXML 2) intensity values for annotated peaks 3) intensity values for un-annotated peaks 4) sample metadata and 5) metadata for the annotated peaks. For each reported peak, information about the analysis mode (reverse phase or hydrophilic interaction chromatography) mass to charge ratio and retention time were collected in the “data_dictionary” tab in the IDSL.CCDB template (SI File 1).

### Processing of the National Health and Nutrition Examination Survey (NHANES) data

Laboratory data for continuous variables were downloaded from the NHANES website (https://wwwn.cdc.gov/nchs/nhanes/search/datapage.aspx?Component=Laboratory) in the SAS export format (.XPT). Variables that reflected a chemical entity were used for calculating the inter-chemical correlation data (Table S1). Data files were imported in the R programming language and merged using the SEQN number as the linking identifier. NHANES data were used for computing correlation statistics without any transformation, normalization and scaling. Survey design weights do not affect the inter-chemical correlations so they were not taken into account.

### Processing of datasets generated by Metabolon Inc. platform

Metabolomics datasets generated by the Metabolon Inc. company available in the supplementary section of a published article^65^ or via metabolomics repositories were included in IDSL.CCDB. The company provides datasets with up to 2,000 high-confidence chemicals reported for blood and urine specimens. For some studies, raw LC-HRMS data are also available through the EBI MetaboLights repository^75^. If these data were not scaled or normalized, we applied a log2 transformation before computing the correlation. For the IDSL.CCDB input format, only metabolite names reported in the table were used in the “data dictionary” tab of the IDSL.CCDB format (SI File 1).

### MS-DIAL data processing

For the MTBLS1684 study, a comprehensive untargeted data matrix was generated using the MS-DIAL software. Parameters were described in the previous report^21^. We applied a log2 transformation before computing the correlation statistics for this study.

### Correlation calculation

The Pearson correlation coefficient was used for computing a pair-wise correlation among reported peaks within each study using the cor function available in the WGCNA R package^76^. A correlation between two intensity vectors was computed only if they had at least 10% non-zero values. We did not compute any p-values for the correlation statistics given that our goal was to create a database of inter-chemical correlations, not to find a biomarker of phenotype. Therefore, the application of a false discovery rate correction was not required. If p-values were computed, they would be expected to be extremely small considering the large sample sizes of the selected studies.

### IDSL.CCDB indexing

For each selected study, a unique name directory was created in a webserver’s filesystem and the pairwise correlation data were saved inside corresponding directory. For each compound, a vector of correlation against all other chemicals in the study were computed and then stored in the file system. For the naming convention, a distinct study-specific identifier was assigned to each reported chemical. Linux operating system Ubuntu 20.04 was used for the webserver.

### Online interface

The online front interface was developed using the AngularJS 1.5 javascript framework and bootstrap. Figure 2 shows a screen shot of the online interface for an untargeted metabolomics study. On the backend, a nginx proxy server was used to route the web requests to the data indexed in the IDSL.CCDB. The opencpu framework in R was used as a middleware to process each web request. For biomonitoring (NHANES), Metabolon Inc’s datasets and untargeted full-scan datasets, three separate types of web-interfaces were developed. For visualizing the correlation data online, Vis.JS javascript library was utilized. If there are more than 100 hits that pass the correlation threshold only the first hundred hits are visualized in the compound centric network and full data were provided as Cytoscape network file.

**Figure 2.**
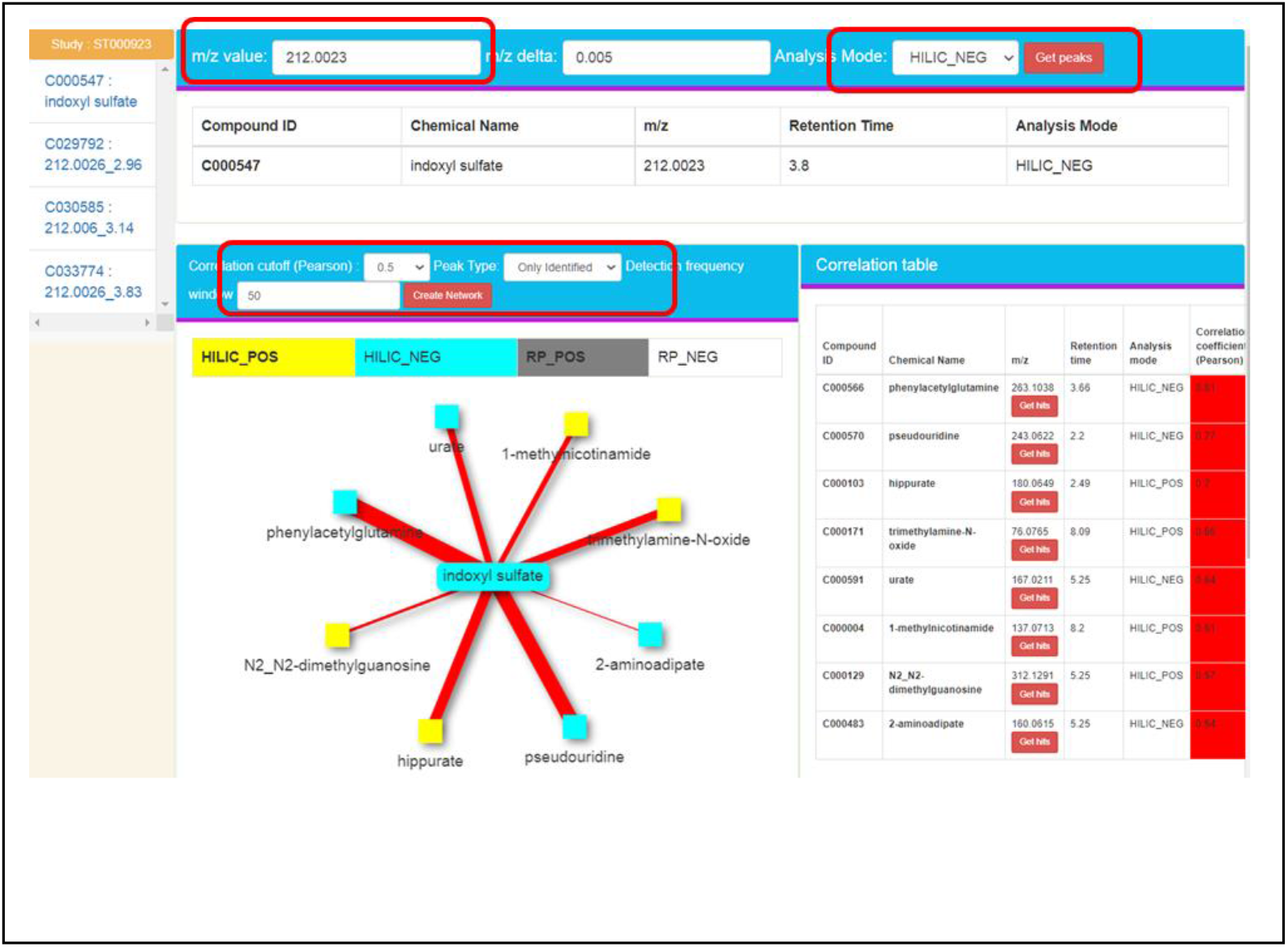
IDSL.CCDB online interface for an untargeted metabolomics study (ST00923). The web interface is available at https://chemcor.idsl.site/originaldata/allpeaks/#?studyid=ST000923

### Querying the IDSL.CCDB

For each study, a specific web-address was created (Table S2). For NHANES data, the query parameter is a variable identifier provided in the Table S1. For Metabolon Inc’s datasets, chemical names were utilized. For full-scan untargeted datasets, m/z with a mass tolerance was used to retrieve the matched peaks in the database. To obtain putative annotation hits, m/z values were matched against a list of compounds that have been associated with a published paper.

### Chemical Similarity Enrichment (ChemRICH) Analysis

ChemRICH is a database independent and *p*-value distribution-based approach to rank the chemical sets that are associated with an exposure^77^. As an example, cor.test function in R was used to obtain *p*-values and estimates for the correlation between Perfluorooctanoic acid (PFOA) intensities and other chemicals from the study IDSLCCDB0001^15^. These results and the sub-pathway information made available by the Metabolon Inc’s report were used as an input for the chemical similarity enrichment analysis using the ChemRICH software^77^.

### Data availability

All data and resources are available at www.ccdb.idsl.me site.

## Results

### IDSL.CCDB is a comprehensive database of inter-chemical correlations for human biospecimens

To build a comprehensive database of inter-chemical correlations in human biospecimens, we found three types of chemical analyses that should be covered. These included 1) biomonitoring surveys that have used a targeted analysis for chemical panels 2) metabolomics datasets having structurally annotated peaks 3) untargeted LC/GC-HRMS datasets having primarily unannotated peaks. In the first version of the IDSL.CCDB, 36 studies were included (Table 1). The coverage for specimen types was 29 (blood), 3 (urine), 4 (stool). The number of individual participants was 107,258 for NHANES with 607 laboratory measurement variables. For 18 datasets that were generated by Metabolon Inc, the sample size ranged from 52 to 1,336 with the reported peak count ranging between 517-1989. For 18 full-scan untargeted LC-HRMS studies, the sample size ranged between 51 to 781 with a reported peak count of 459 to 81867, and 9 studies had reported only un-annotated peaks that were referenced using m/z and retention time values. Of the untargeted studies, 14/18 studies had data collected in the reversed phase mode, and 8/26 had data collected in the HILIC mode, and 5/26 had data collected in both RP and HILIC modes. To update the database, we plan to regularly screen publicly available datasets in the NIH MetabolomicsWorkench, EBI MetaboLights, GNPS-Massive and consortium/cohort specific repositories and supplementary tables for published papers and include the relevant studies in the IDSL.CCDB database. By covering three types of chemical measurement datasets, IDSL.CCDB can provide unique opportunities to not only learn about the biological relationships among metabolites, but also prioritize chemicals that are yet to be annotated in untargeted LC/HRMS datasets.

### A large number of inter-chemical correlations were observed in the catalogued studies

To populate the database, pair-wise correlations among reported chemicals were computed for each selected study. A computational pipeline has been established for an efficient indexing of a new dataset in the database. For that, a minimal level of manual curation was needed to prepare the dataset in the required format (SI File 1). We investigated the prevalence of strong inter-chemical correlations across the catalogued studies. A total of 176.6 million inter-chemical correlations across the studies passed a threshold of 0.6 Pearson coefficient, indicating the large-scale and magnitude of strong correlation patterns that exists among chemical compounds present in human biospecimens (Figure 3). More of these correlations were observed for untargeted datasets which had thousands of mostly unannotated peaks. We noticed that endogenous compounds tend to show a higher number of significant correlations in comparison to exogenous and xenobiotic compounds (Figure S1). This suggested that at a lower correlation threshold level, we can capture new relationships among chemicals that would otherwise be missed if the correlation data is visualized as a network graph created using a stringent threshold.

**Figure 3.**
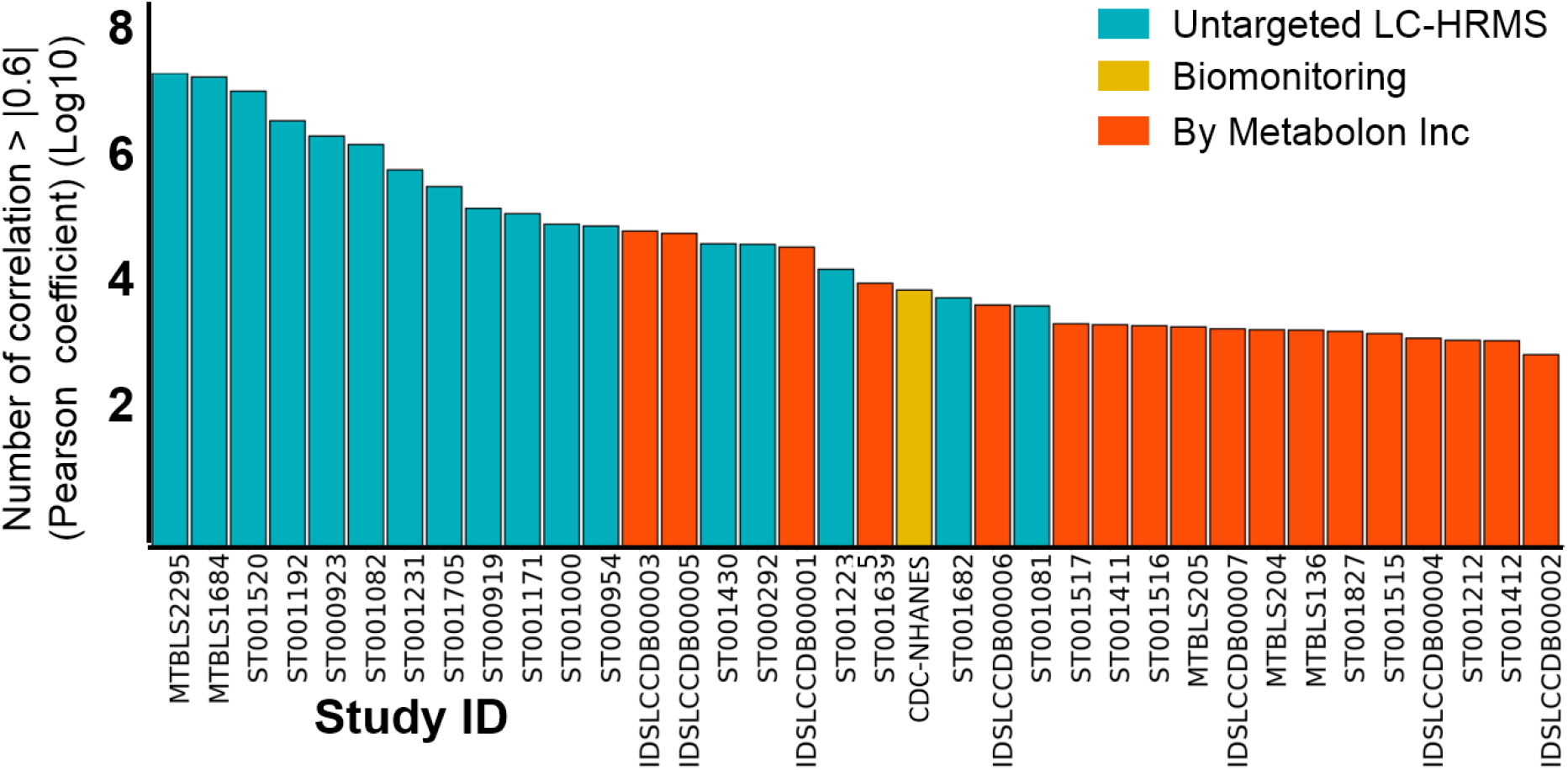
Prevalence of strong inter-chemical correlations across 36 studies in the IDSL.CCDB. These are unique correlations. See the Table 1 for the description of each study and number of compounds.

For example, by a Pearson coefficient cutoff of 0.4, we have noticed a relationship among blood glucose and acyl-choline lipids (Figure S2) in the study MTBL136^58^ which will be missed on a cutoff of 0.6. This association has been linked with energy disturbance and implicated in diabetes and chronic fatigue disease related studies. This underscores the need to access the correlation data in a flexible and interactive approach so we can capture both the known and novel types of functional and biological relationships among reported chemicals.

### Dataset type specific web-interfaces provided access to correlation data for both annotated and unannotated compounds

Because a large number of inter-chemical correlations were observed in the selected studies, it was not practical to visualize them in Cytoscape network visualization^78^ or any other network graph visualization software unless the network graph is created using very stringent correlation thresholds, which will likely miss biological insights. Therefore, we stored all the correlation data for each compound from each study in a web-server’s file system. This allows us to readily load the correlation vector in the computer memory without the need to re-calculate them and enabled a faster response time for the online visualization. Also, a compound centric view was found to be a cleaner, readable and meaningful visualization than creating a network graph of all compounds reported in a study. It enables a focused investigation of a single compound and its chemical and biochemical relationships with other chemicals in a study. A network-based visualization highlighted a compound centric view of inter-chemical correlations (Figure 2), which can be updated by different correlation thresholds. Three types of web interfaces were developed to provide a tailored access to biomonitoring, annotated peaks and unannotated data in metabolomics and exposomics assays (Figure S3–5). These interfaces enabled queries by chemical names, CAS numbers, NHANES identifiers and mass to charge (m/z) ratio. For untargeted assays, data from different analysis modes were stacked which allowed to find peaks from the same compound in two analysis modes such as a ESI positive and negative or HILIC (+) or RP (+) (Figure S6). Network data were also provided as Cytoscape network files to enable additional visualization strategies. These simple and flexible web-interfaces allowed a seamless and interactive access to the inter-chemical correlation data for a chemical from a study.

### Compounds from a chemical class correlated strongly with each other in the NHANES biomonitoring dataset

First, we asked if compounds from a known chemical class correlate with each other and can be retrieved by querying a single chemical. We have observed that chemicals from well-recognized environmental exposures PCB, PFC and PAH groups indeed correlated with a representative chemical from these classes (Figure 4). This probably suggested a common source of exposure for these chemicals. When cotinine, a biomarker of tobacco smoke was queried, it retrieved many other tobacco smoke related chemicals, providing a quick overview of biomarkers of smoke exposures.

**Figure 4.**
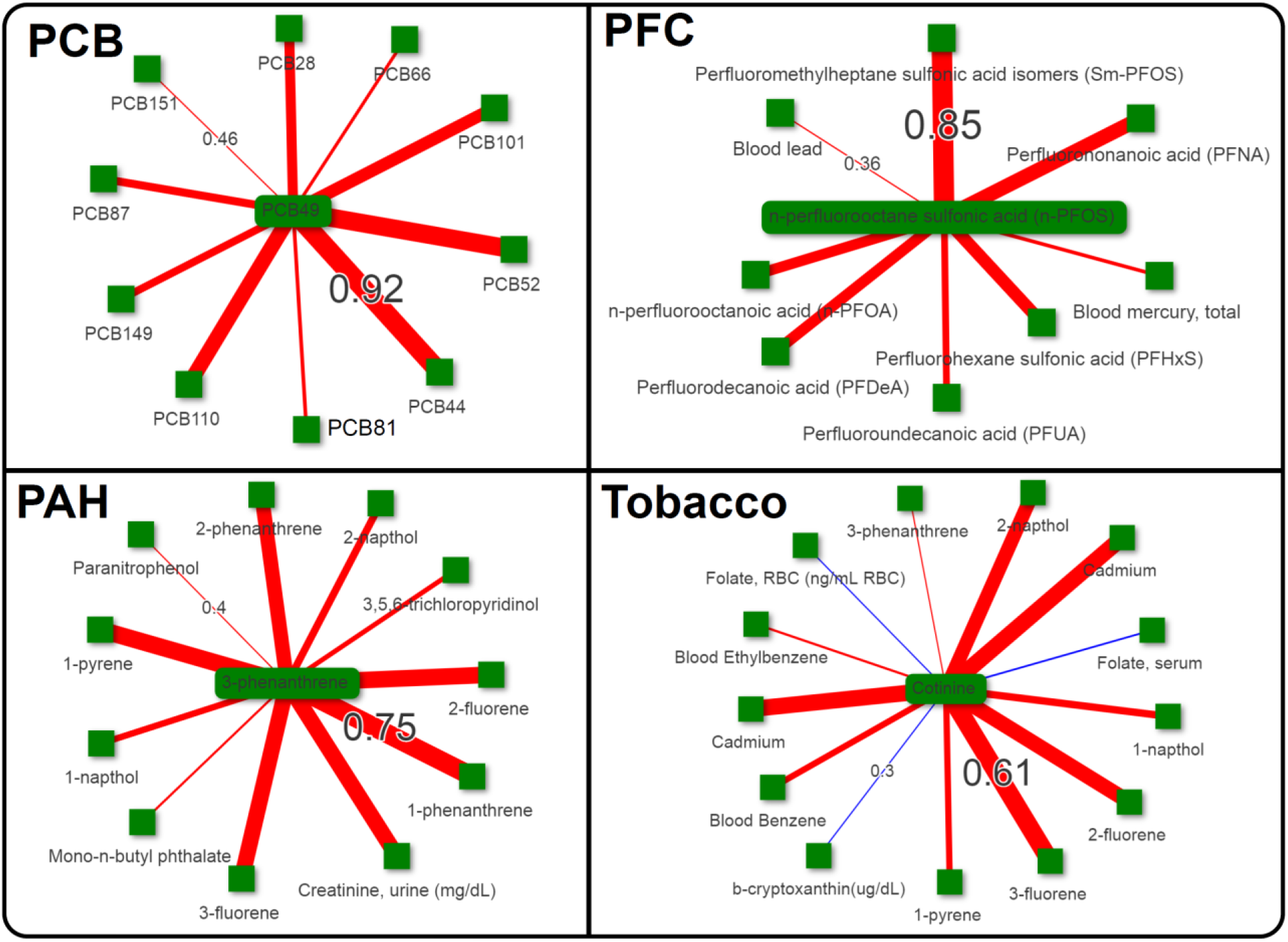
Correlations among chemicals within a class or having same source origin in the NHANES dataset. Thickness of lines indicate the correlation strength. The correlation cutoff was 0.3 for PCB, PFC and Tobacco compounds, and 0.4 for PAHs. PCB - Polychlorinated Biphenyls; PAH-polyaromatic hydrocarbons, PFC - Perfluorinated compounds. Online network can be accessed at the site - https://chemcor.idsl.site/originaldata/biomonitoring/#?studyid=NHANES Only minimum and maximum correlations are shown on the edges for clarity. Thickness of edges are not comparable in two network figures.

This compound-centric retrieval of inter-chemical correlations in the NHANES biomonitoring dataset suggested that chemical exposures with similar structure and origin correlates strongly with each other. Additionally, IDSL.CCDB database can be utilized to prioritized chemical exposures for new biomonitoring studies using targeted analysis.

### Stronger correlations among compounds belonging to a chemical class in metabolomics datasets

Next, we investigated if endogenous metabolites from a chemical class correlate with each other in a metabolomics dataset. We queried a ubiquitous endogenous blood metabolite, C-16 carnitine and retrieved its neighbors in the IDSLCCDB00007 study. At the Pearson correlation cutoff of 0.6, we retrieve mostly other saturated and unsaturated carnitines (Figure 5). However, at the 0.5 Pearson correlation cutoff, we found that carnitines have biochemical relationships with fatty acids and sphinganine metabolites.

**Figure 5.**
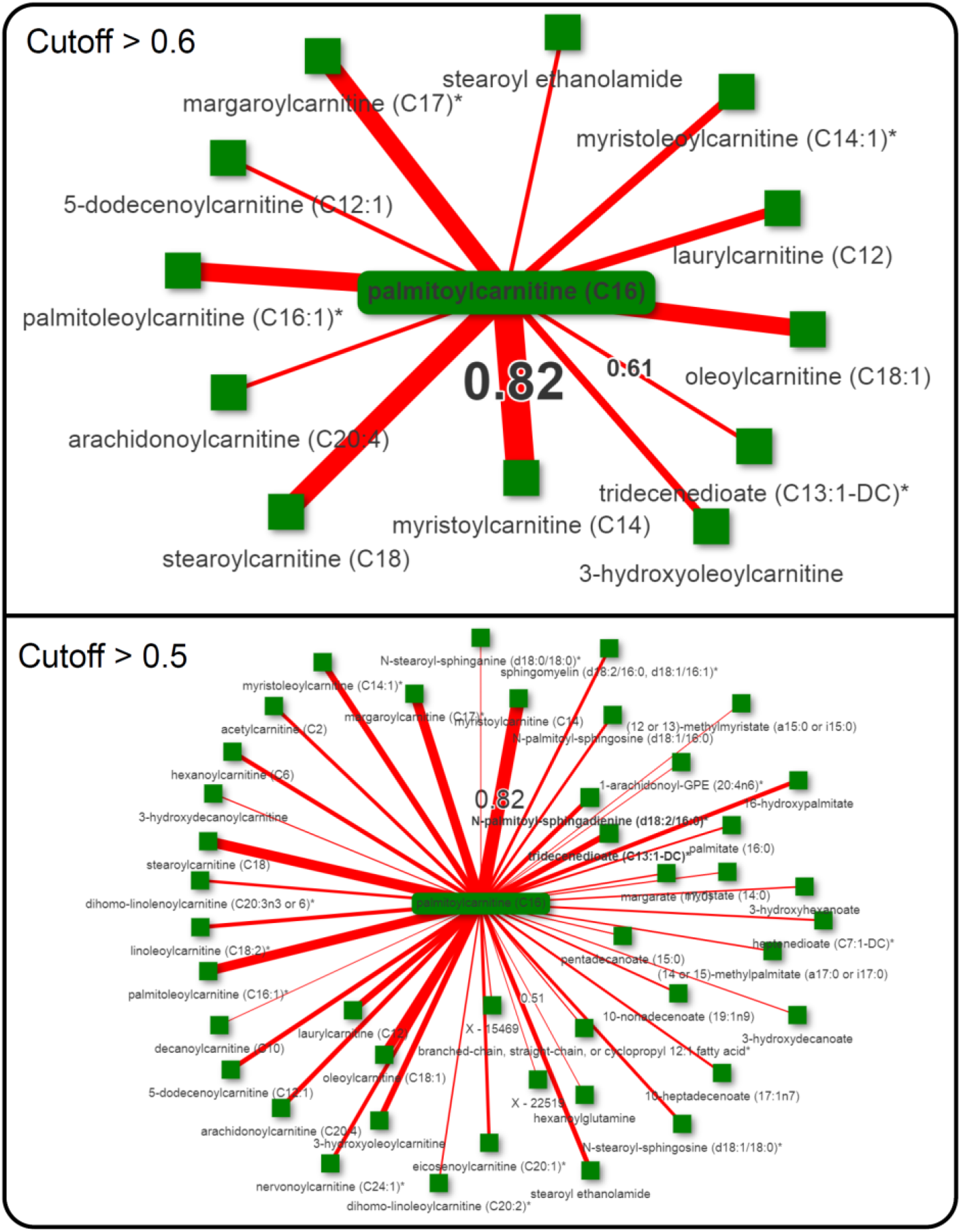
Compounds correlation with CAR 16:0 in the study IDSLCCDB00007. Only minimum and maximum correlations are shown on the edges for clarity. Thickness of edges are not comparable in two network figures.

We learned that structurally similar compounds from an endogenous chemical class can have a very high correlation coefficient among them, suggesting an enzyme activity that can react on any member of a chemical class, for instance the carnitine palmitoyltransferase I enzyme. As the Pearson correlation cutoff was lowered, we found long-distance biochemical relationships suggesting different chemical classes that may belong to a metabolic pathway, for instance, fatty acids and acyl-carnitines. It also highlighted the unidentified metabolites in the Metabolon’s report which may belong to the acyl-carnitine chemical class. Overall, by modifying the correlation cutoff, the IDSL.CCDB interface enables retrieval of short and long-distance biochemical relationships in a metabolic network around a single chemical. This can be used for hypothesizing novel biochemical relationships in untargeted metabolomics datasets.

### Products of xenobiotic metabolism

Next, we checked if metabolites of a xenobiotic compound correlate with the parent compound’s levels. In the NHANES biomonitoring survey, several metabolites of caffeine strongly correlated with caffeine levels (Figure 6, upper panel). The same pattern was found in a metabolomics dataset (Figure S7). Similarly, metabolites of mono-n-butyl phthalate (MnBP), a commonly used plasticizer correlated with structurally and metabolically related chemicals. MnBP also correlated with other phthalate molecules (Figure 6 lower panel), indicating common exposure sources. It was expected that people exposed to dibutyl phthalates will excrete MnBP and mono-isobutyl phthalate in their urine (ref CDC). For acetaminophen, a commonly used over the counter pain-reliever drug, its sulfate metabolite was found to be correlating with other acetaminophen metabolites (Figure S8).

**Figure 6.**
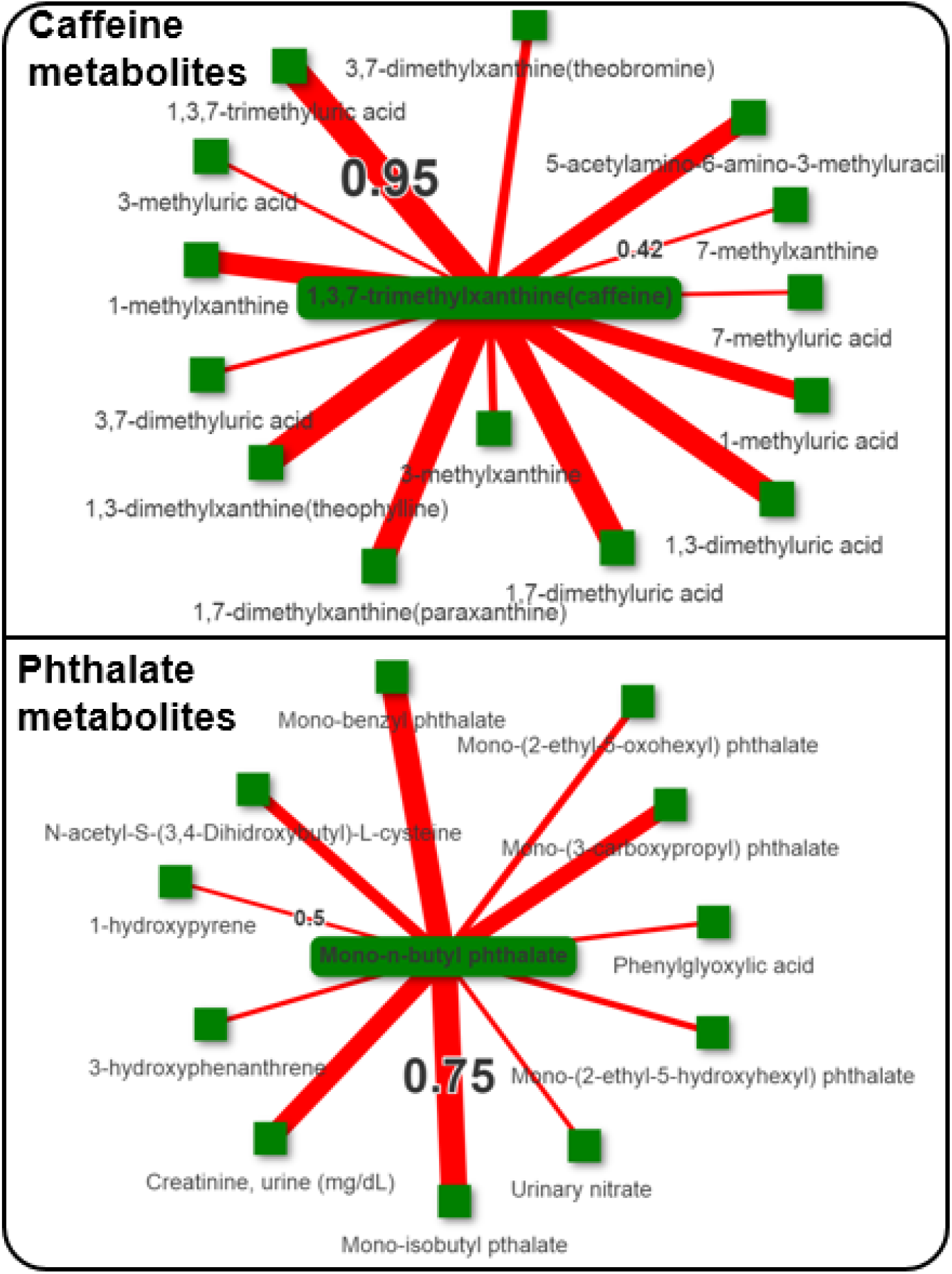
Caffeine and phthalate metabolites in the NHANES survey data. Variable id URXMBP_PHTHTE_D (year 2005-2006) was used for mono-n-butyl phthalate (MnBP). Variable id URXMX7_CAFE_H (year 2013-2014) was used for caffeine. Label on the edges show the Pearson coefficient. Only minimum and maximum correlations are shown on the edges for clarity. Thickness of edges are not comparable in two network figures.

### Putative annotation of peaks in untargeted data by correlation patterns

So far, we have learned from the NHANES and other high quality metabolomics dataset that chemicals within a chemical class or having the same origin or similar pathway tends to show strong correlations. Relying on this information, we explore the untargeted metabolomics datasets to test if m/z values for chemicals from a chemical class show inter-chemical correlations. To test this, we have queried the m/z value 498.9291 for the M-H adduct of perfluorooctanesulfonic acid (PFOS) in reverse phase chromatography data for the ST001430 study. It retrieved three other chemicals on in the correlation cutoff of 0.3, which matched to the M-H adducts for other common PFCs - PFOA and PFHxS (Figure 7). In another untargeted study ST001231, we found that PFOS correlated with many more PFCs compounds (Figure 7).

**Figure 7:**
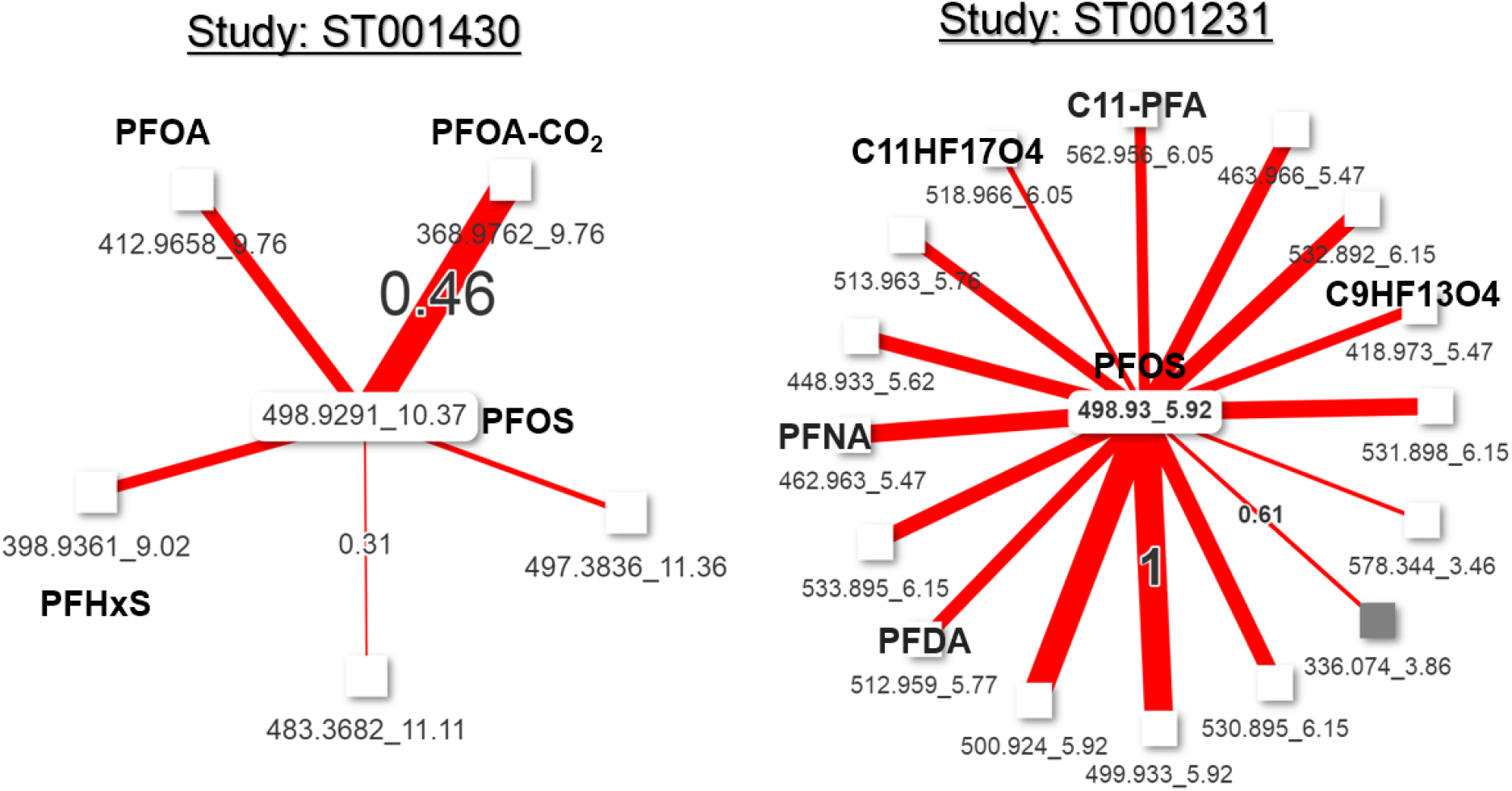
Inter-chemical correlation among PFCs in the untargeted metabolomics datasets. Correlation threshold for ST001430 was 0.3 and for 0.6 for ST001231. Only minimum and maximum correlations are shown on the edges for clarity. Thickness of edges are not comparable in two network figures.

### Metabolic effect of a hazardous chemical - PFOA

Finally, we asked if we could utilize the inter-chemical correlation data to understand metabolic effects of a chemical exposure. Perfluorochemicals (PFCs) are concerning chemicals for public health. They are exclusively synthetic and accumulate in human body overtime. The ubiquitous exposures to them have been under high priority investigations since they may have contributed to the etiology of a range of chronic diseases. Endogenous metabolites that correlate with PFCs exposures may reflect the biological response to these hazardous chemicals. In several of Metabolon Inc’s reports, Perfluorooctanoic acid (PFOA) peak was annotated and found to be correlated with many chemicals when we indexed these reports in the IDSL.CCDB.

Many metabolites that correlated with PFOA levels may belonged to the same pathway or chemical class. Identifying these chemical sets can assist in understanding the systematic metabolic effect of PFOA exposure which can span over multiple metabolic pathways (Figure 8). Therefore, we have utilized ChemRICH analysis^77^ to identify the PFOA associated chemical sets, which suggested that PFOA exposure has a negative effect on most of the lipid sets except triglycerides^79, 80^. PFOA exposure may have also induced the amino acid and tocopherol metabolism pathways. This analysis highlighted that IDSL.CCDB correlation data can also be used for investigating the metabolic hazardous effect of a chemical exposure of public health concern using a chemical set analysis approach.

**Figure 8.**
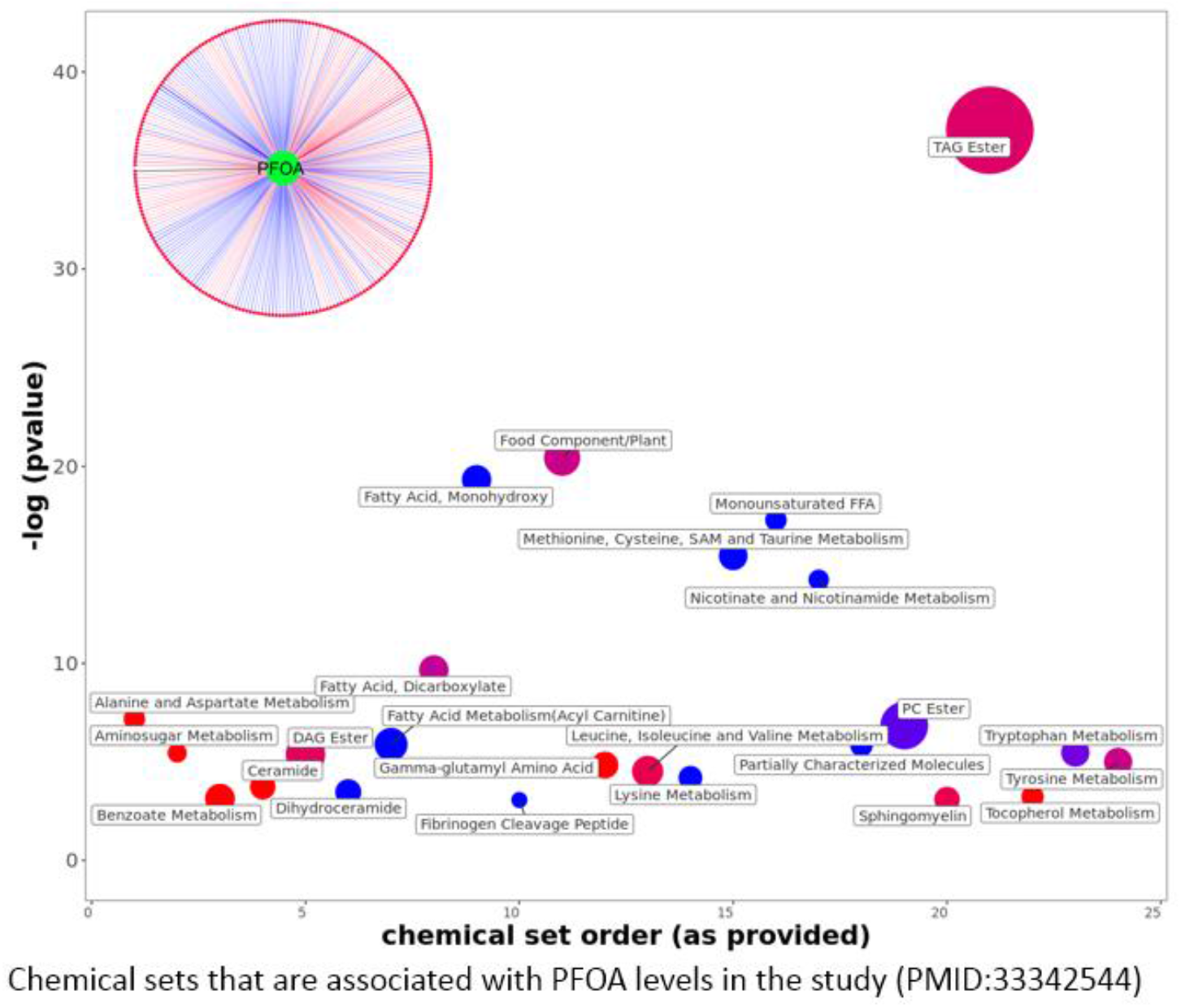
Chemical similarity enrichment analysis of PFOA and its correlation with other metabolites in a report generated by Metabolon Inc. The circular diagram shows individual chemicals that correlated with PFOA levels.

## Discussion

Inter-chemical correlations in biomonitoring, metabolomics and exposomics datasets is a useful source of information to expand our understanding about the relationships between different metabolites, metabolic pathways and the chemical exposures. There is a need to systematicaly catalogue and preserve these correlation patterns in a database to support a range of meaninful queries. In this paper, we have presented the IDSL.CCDB database which aims to build a catalogue of inter-chemical correlation in chemical measurement datasets and then provide users access to the correlation data using a web-interface. As of January 2022, the database includes data from from 36 studies covered. We plan to regularly screen literature as well as metabolomics and exposomics repositories to identify additional studies that can be catalogued in the IDSL.CCDB. The database currently only hosts studies related to human specimens, however given the generic nature of the catalogued data and indexing pipelines, it will be able to incorporate studies of other species or sources as long as data are provided in the required format. We foresee a regular use of the database in the field of metabolomics and exposomics to explore about the biochemical and chemical relationships around a chemical that has been prioritized by a researcher using statistical or by text mining approaches. We believe the IDSL.CCDB will be a core database resource in these fields where the interpretation of multi-analyte datasets remains a major challenge.

A large number of significant inter-chemical correlations are ubiquitously observed in these core datasets. An obvious question is “what are the reasons behind these correlations”? At present this is a challenging question because these correlations can be interpreted only in the context of known exposure sources, biochemical absorption pathways and transformation reactions^81^. With time, and additional cataloguing of the exposome, the reasons behind these correlations will become more evident. Pathway-centric approaches do not cover many high-priority exposome chemicals, including their chemical classes, source origin and transformation products. The IDSL.CCDB is designed to address this issue by curating and interpreting inter-chemical correlations in exposomics and metabolomics core datasets while integrating information about functional and structural relationships among chemicals.

In the transcriptomics field, gene correlation or co-expression databases^33, 82^ have been developed for multiple species and disease conditions. These databases allow the identification of gene function(s) based on the similarity between two gene’s expression levels. They have shown that the similarity in expression levels reflect a shared function or regulation in the genetic networks. IDSL.CCDB is in line with these databases to provide similar resource for chemicals. For the first time, we developed an inter-chemical correlation database to be used for metabolomics and exposomics hypothesis generation and characterization.

Due to the large number of analytes in targeted and untargeted assays, a traditional correlation network graph^83, 84^ using Cytoscape^78^ or similar software would not be meaningful to explore inter-chemical correlation data because the network graphs would be over-crowded requiring a stringent correlation threshold to draw the edges. Instead, we propose to use a single-compound centric network to generate clear and readable networks that are easy to deploy in online interfaces. We suggest that investigators can explore correlation data in this interactive, compound centric way so that novel relationships among chemicals can be readily explored. In this way, IDSL.CCDB fullfills critical gaps in the mining of metabolomics correlation data.

IDSL.CCDB can play a role in peak annotation in untargeted metabolomics, because compounds belonging to the same class, metabolic pathway or source origin tends to correlate with each other. By querying a single compound’s m/z, we will be able to estimate the chemical class or in some cases the exact identity of a peak, although it will be only based on the MS match against a priority list of chemicals from a database. There is a need to develop further tools to utilize the isotope patterns, MS2 spectra to refine the annotation patterns. For full-scan untargeted datasets, m/z with a mass tolerance will be used to retrieve matched peaks in the database. In untargeted chemical analyses, many inter-chemical correlations are often observed due to non-biological causes. They are useful in annotating peaks in the untargeted dataset with isotope information^85^, chemical fragments^86^, and errors during data processing, such as duplicate peaks. These annotations can be transferred to other untargeted studies with many unidentified peaks, with the logic that pairs of the same compound will show similar inter-chemical correlation irrespective of analysis platform. It was shown in the example for PFCs and caffeine metabolites (Figure 8 and S8). It is expected that some of the inter-chemical correlations may not be found across multiple studies or may not have the same strength, which can suggest that the underlying regulatory or source mechanisms are operating differently in two studies. These differences can be considered high priority hypothesis.

Future directions will include enriching the correlation database with more tissue types and clinical outcome data. This will enable us to highlight the biomedical relevance of a compound-centric correlation network that is created for a phenotype or outcome. We will also include publicly available datasets from the HHEAR program and other NIH supported consortiums. We will also add text mining^23^, chemoinformatics and other bioinformatics resources^81^ which can help in interpreting the inter-chemical correlations in an automatic manner. For the untargeted metabolomics dataset, we need to re-generate those datasets using MS-DIAL^87^ or other software which can generate comprehensive data matrices so that the chances of missing any biological insights will be minimal^21^.

## Conclusions

We describe IDSL.CCDB, a new key database in the field of metabolomics and exposomics that provides access to fundamental information on the inter-chemical correlations among chemical signals derived from human specimens. The database has great potential to accelerate learning about the chemical and biochemical relationships among reported chemicals. It can be used for prioritizing chemicals, identifying new hypotheses, interpreting metabolomics datasets, annotating peaks in untargeted metabolomics datasets, and for investigating the metabolic effects of a known chemical exposures. Overall, IDSL.CCDB will start a new wave of database types in the metabolomics and exposomics field that are more interpretive than just a catalogue of information.

## Author contributions

DKB and ST conceptualize the study. DKB and SFB prepare the data and conducted data analysis. PM and DKB designed the web-interfaces and database architecture. All authors contributed to the manuscript.

## Supplementary information

**Table S1:**
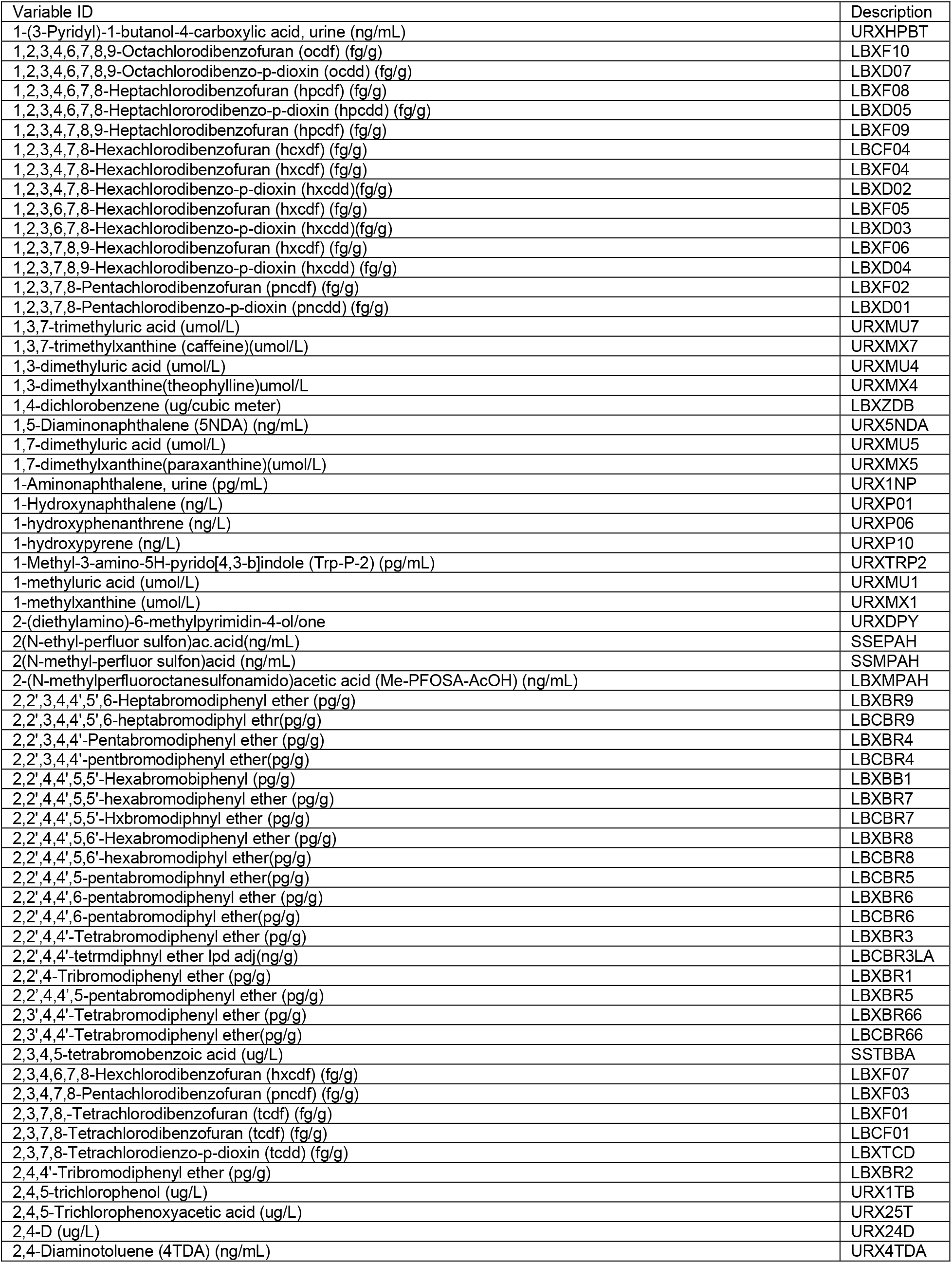
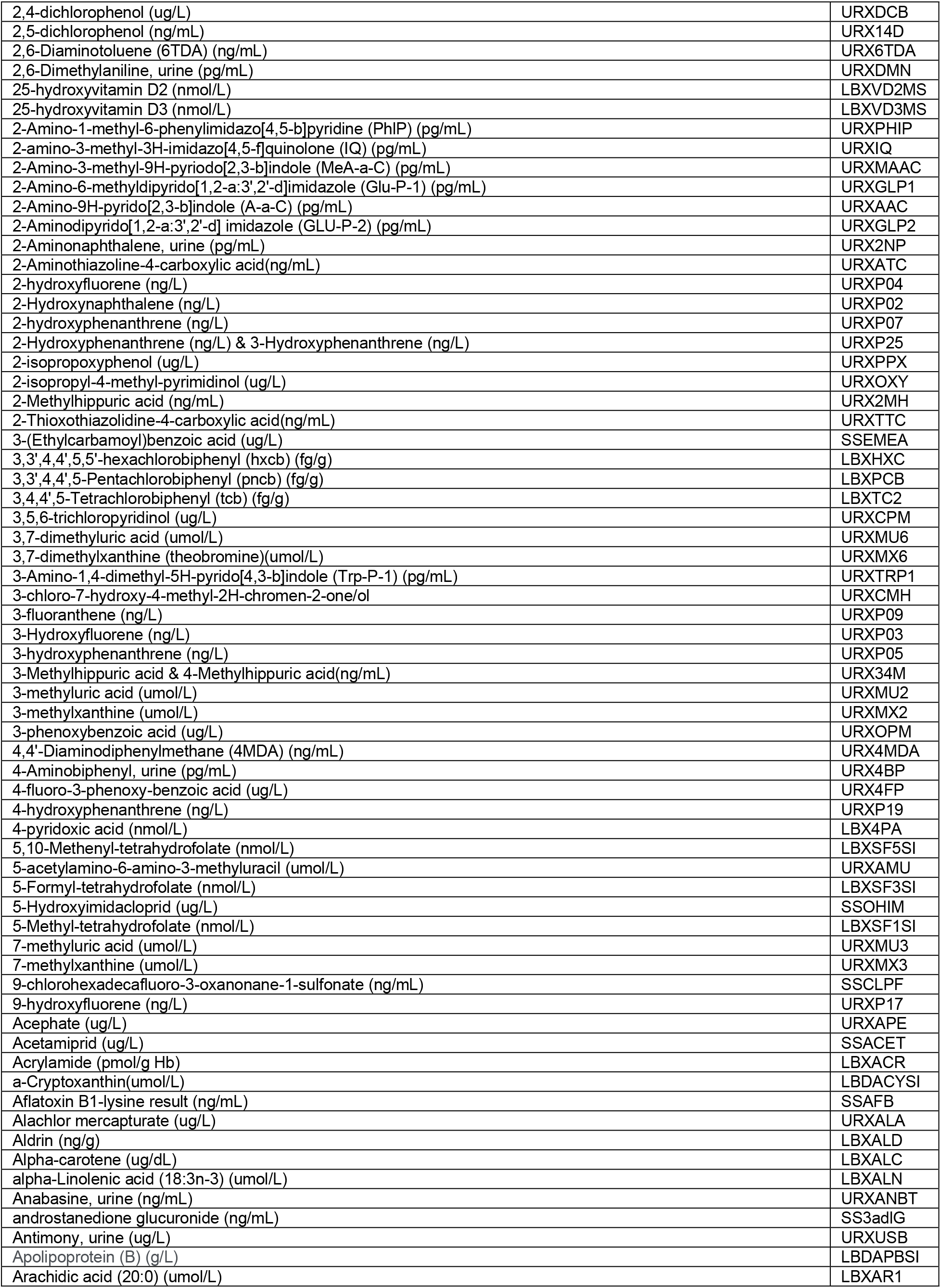
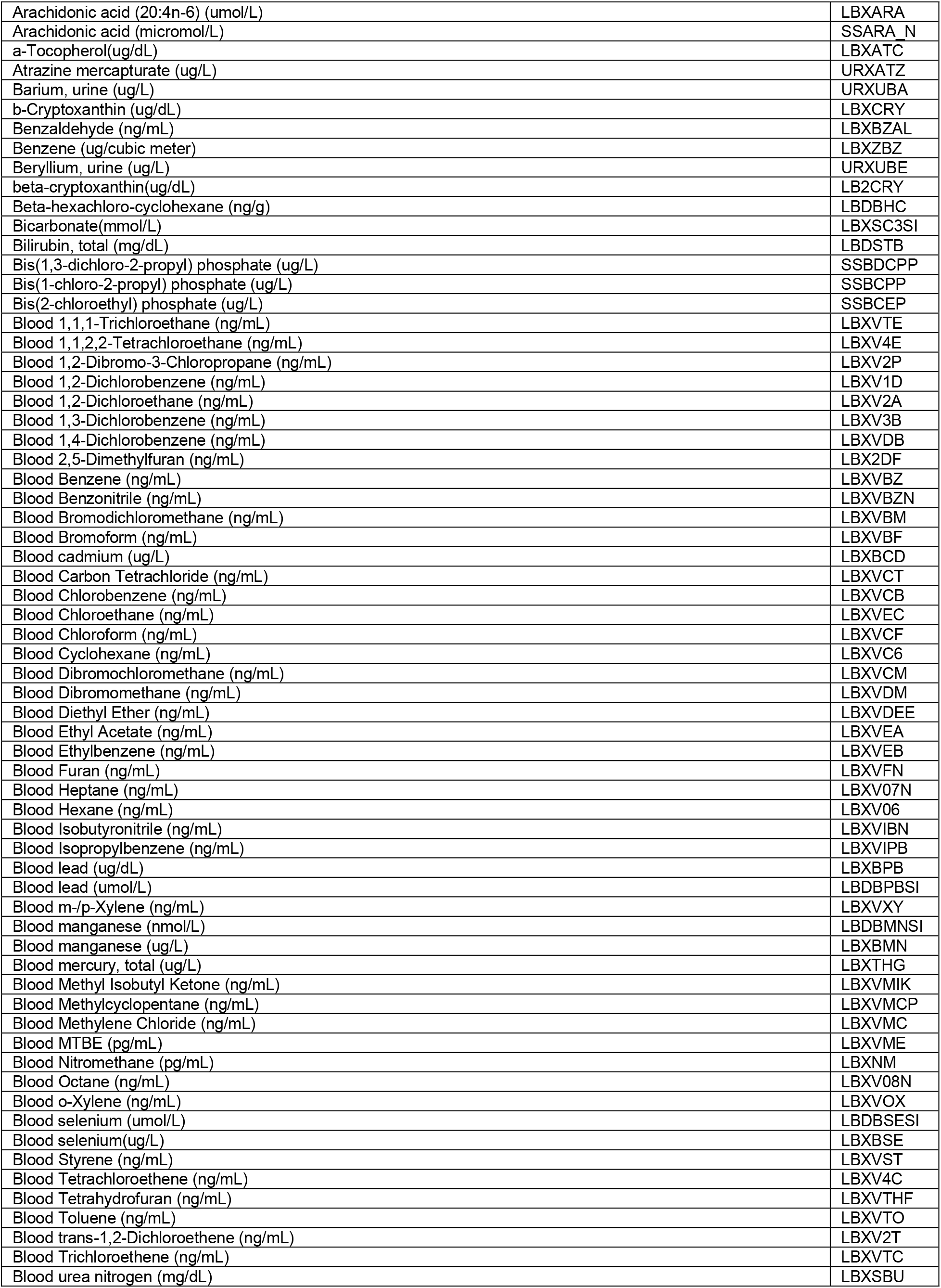
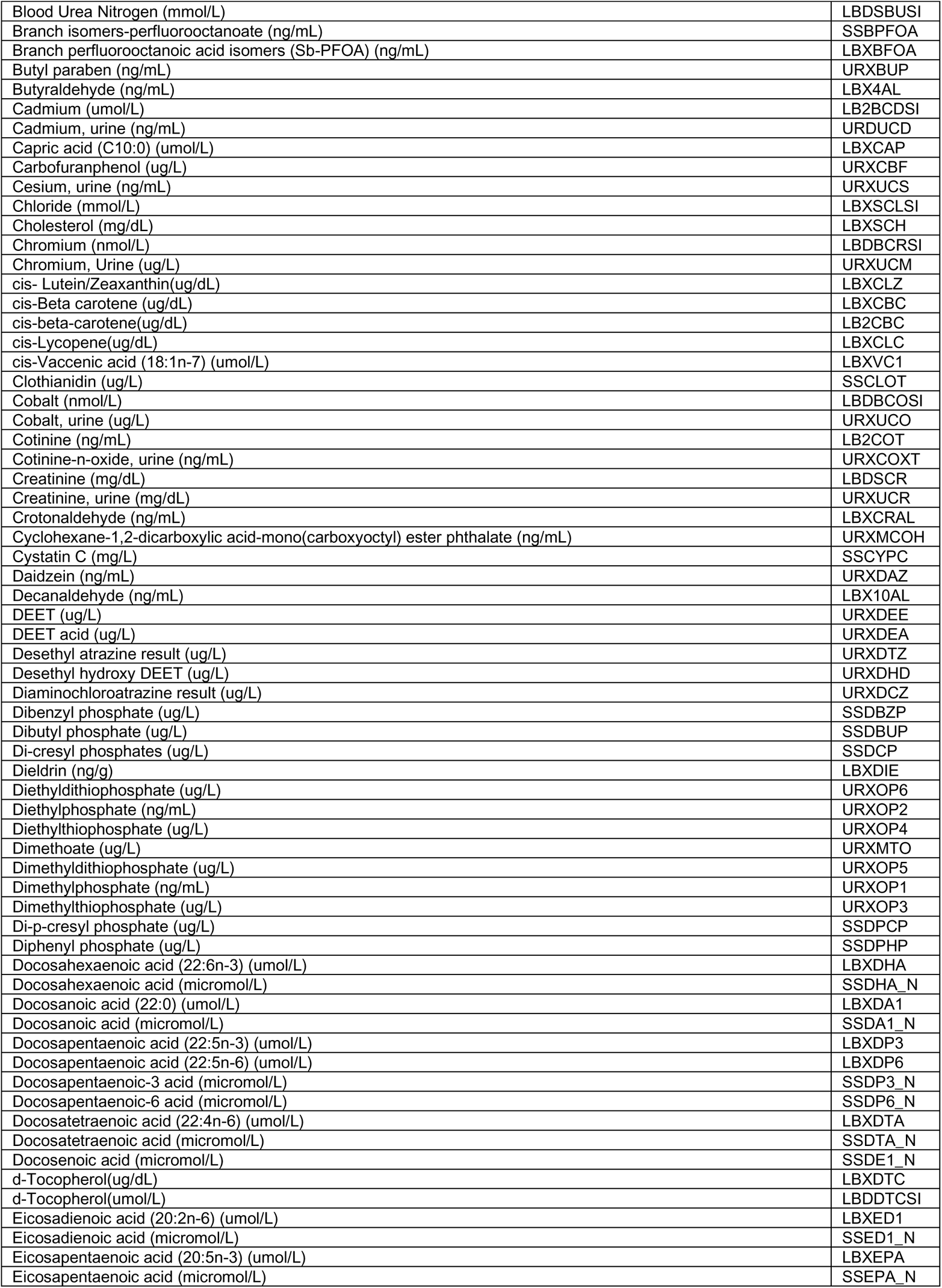
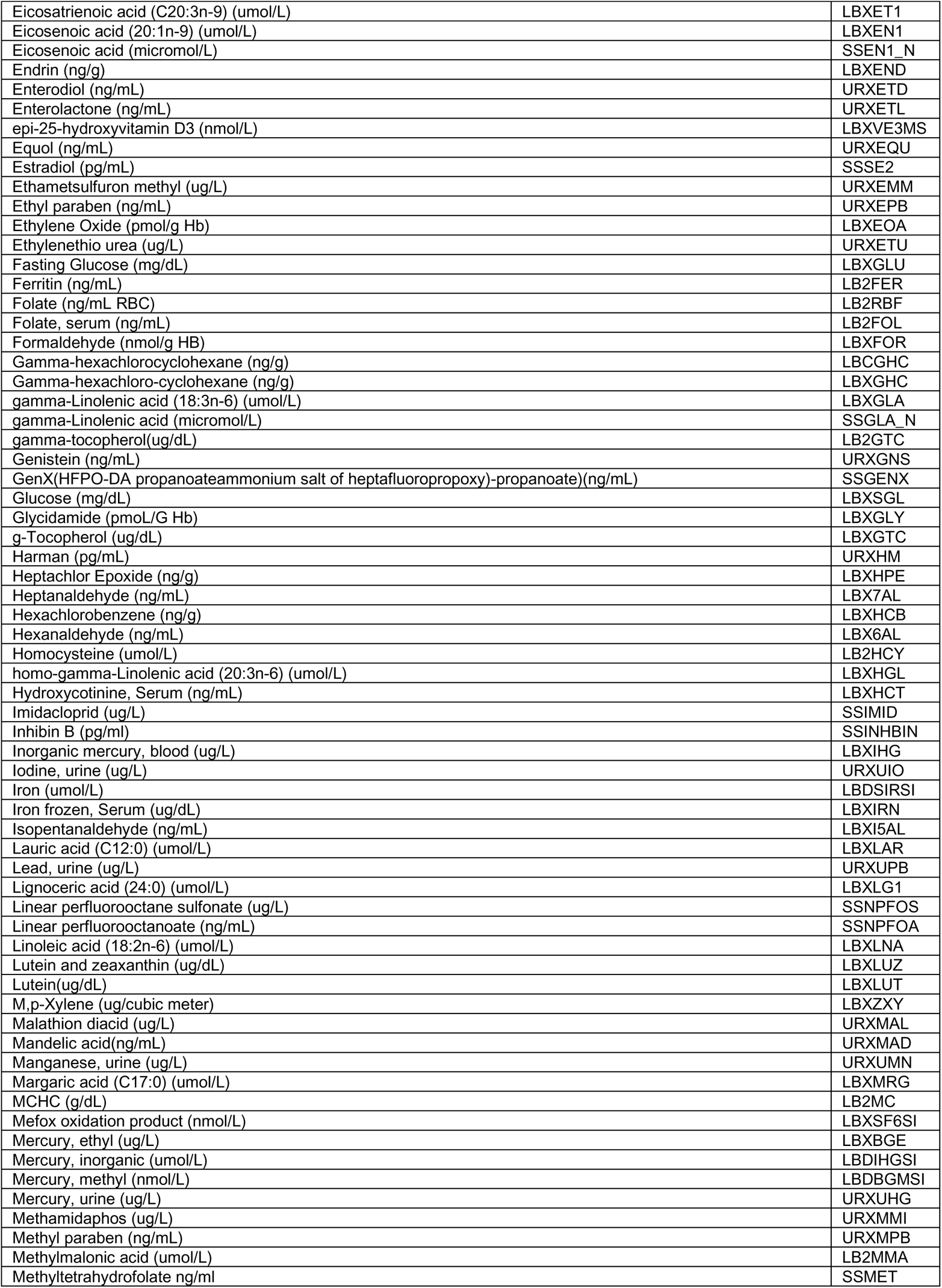
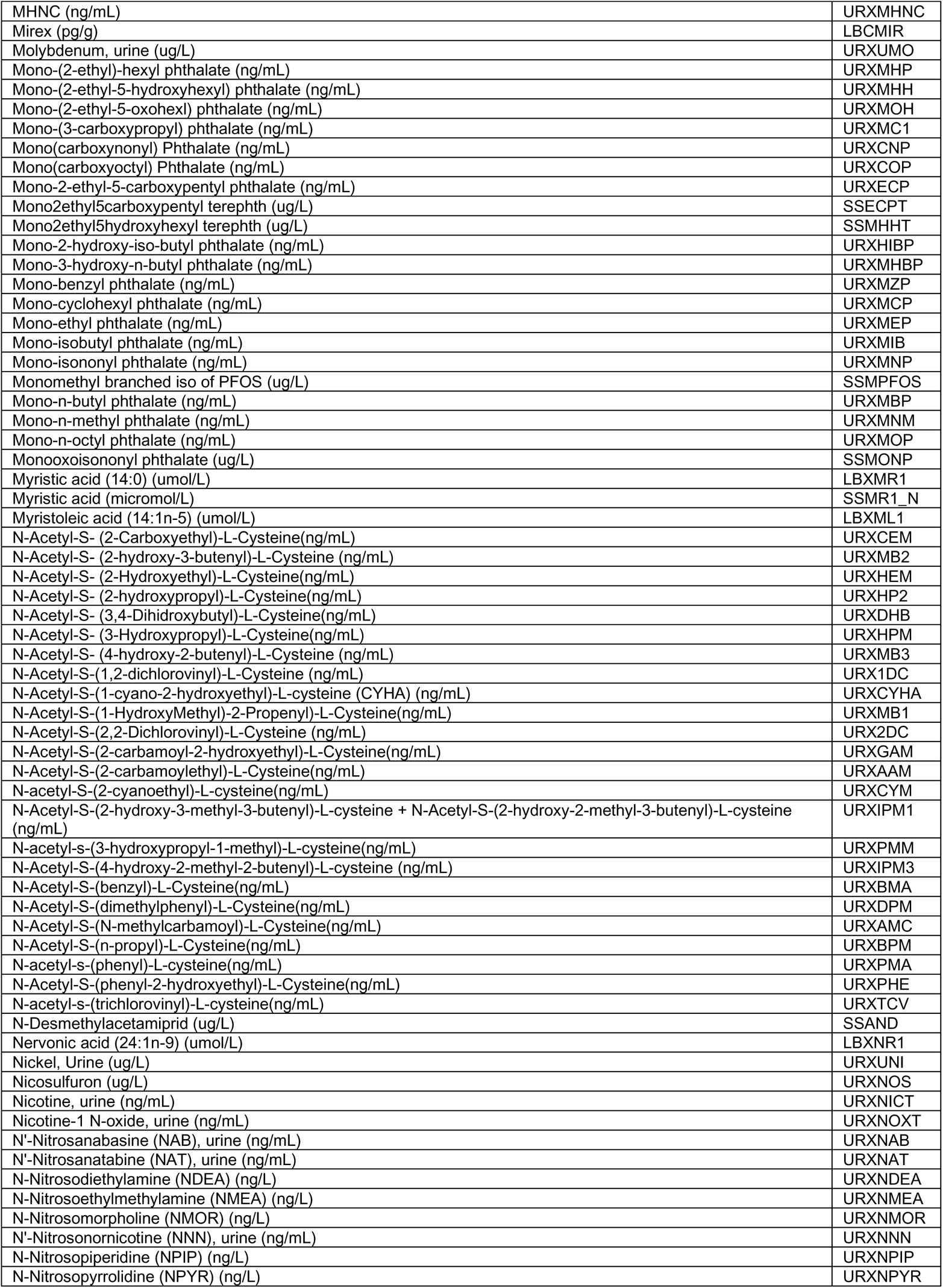
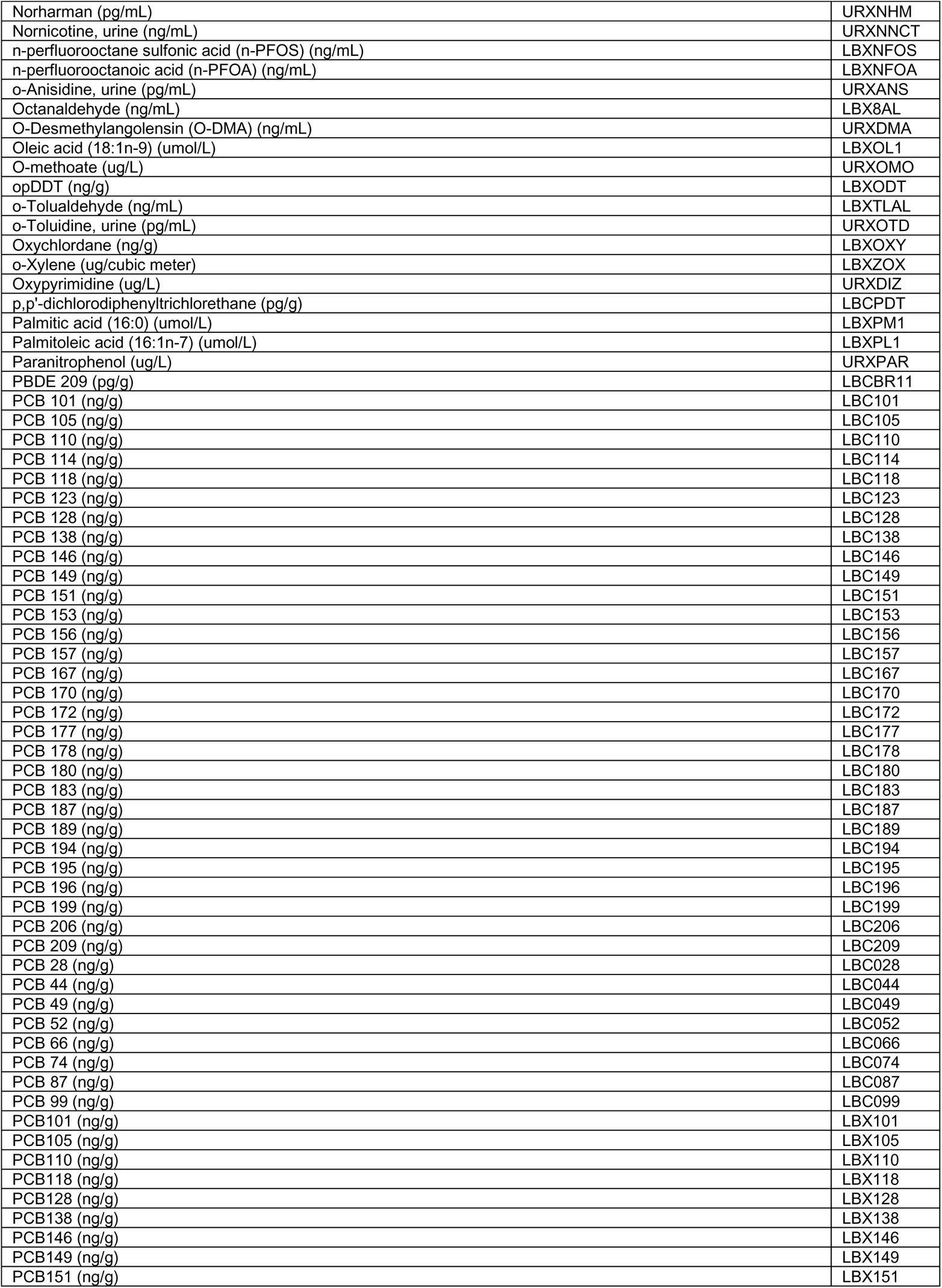
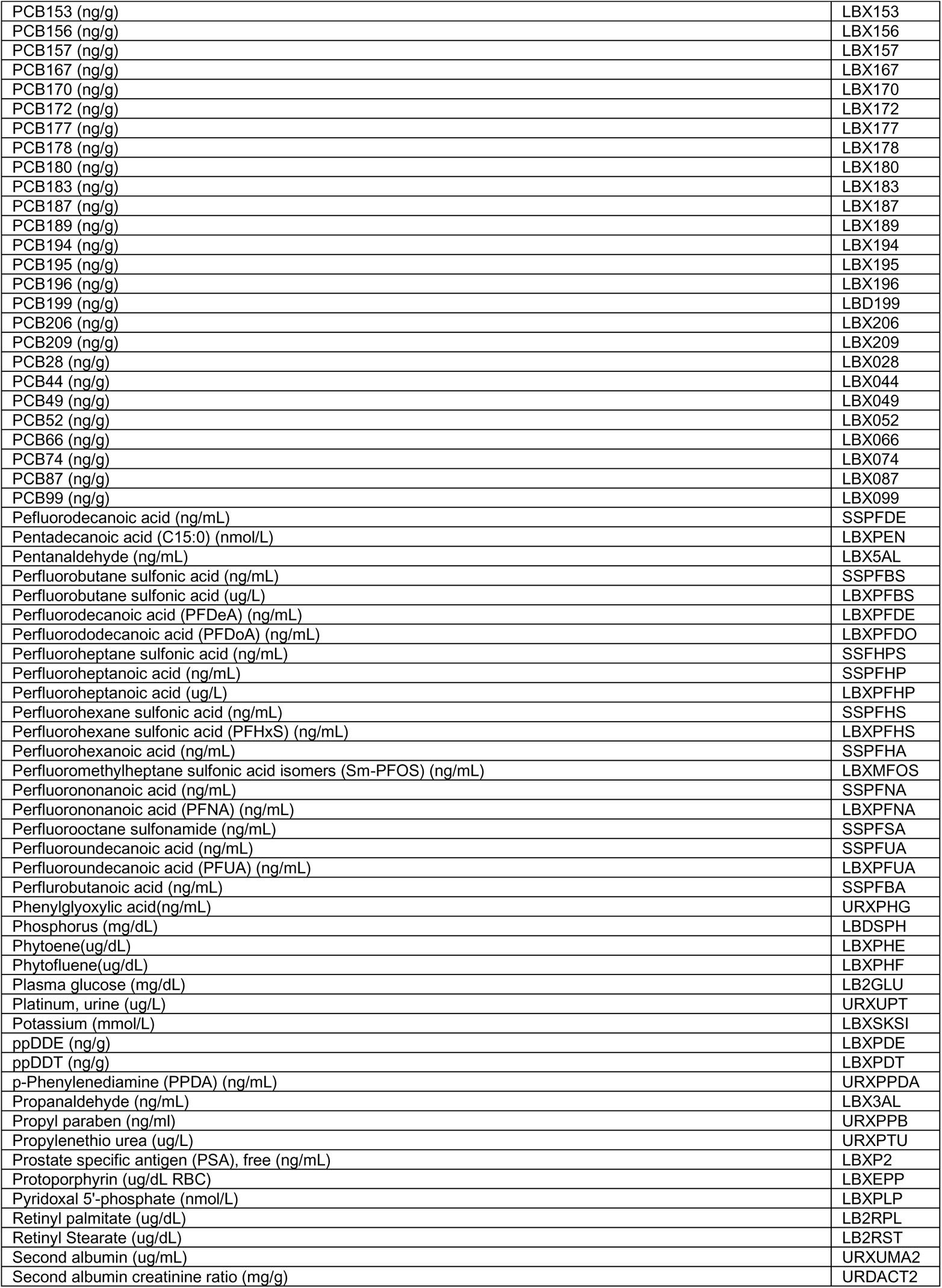
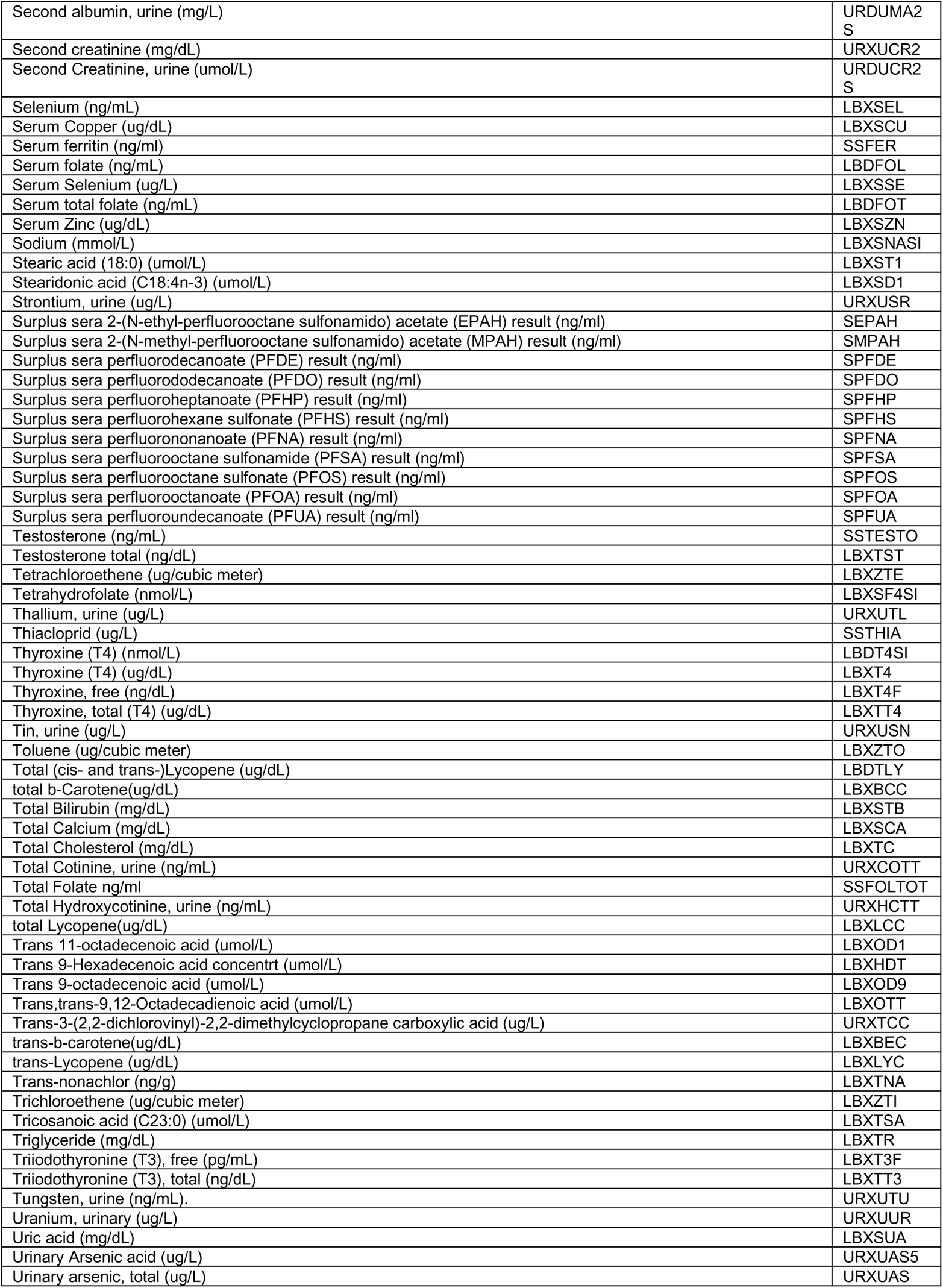
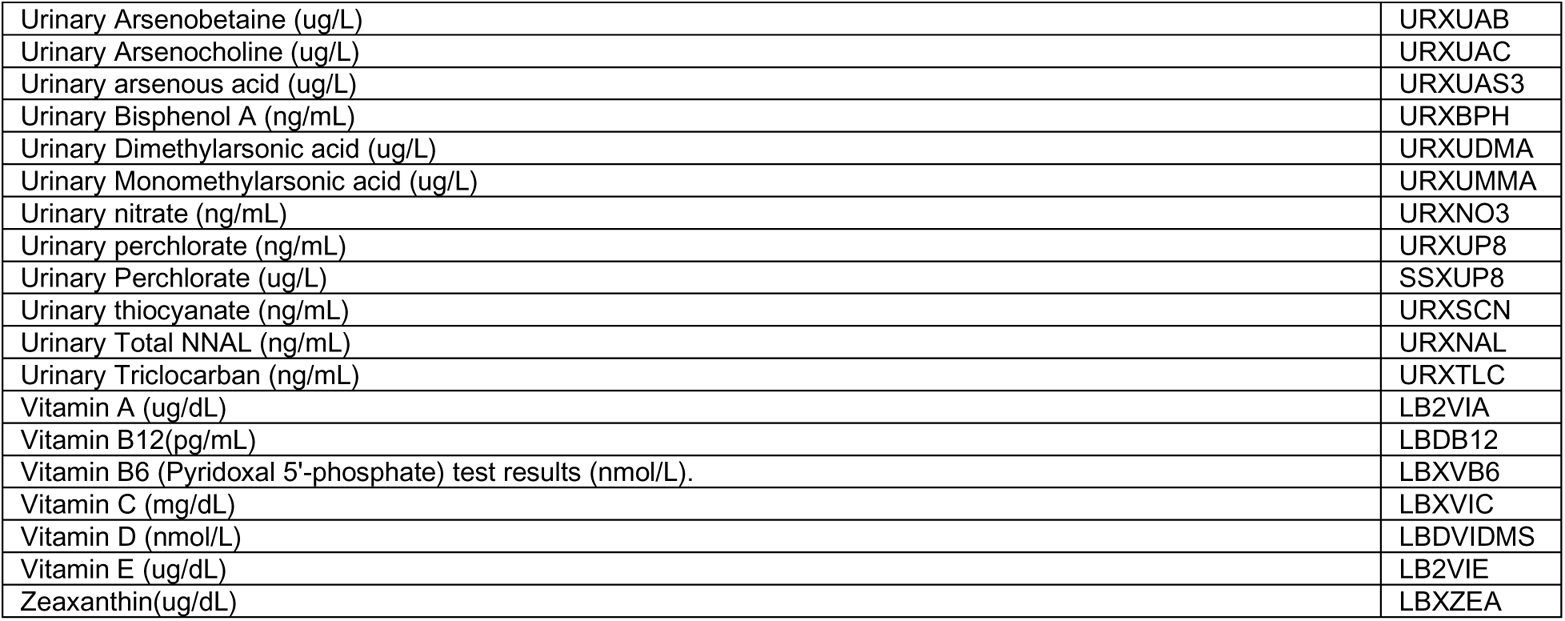
NHANES variable list that can be queried in the IDSL.CCDB database.

**Table S2:**
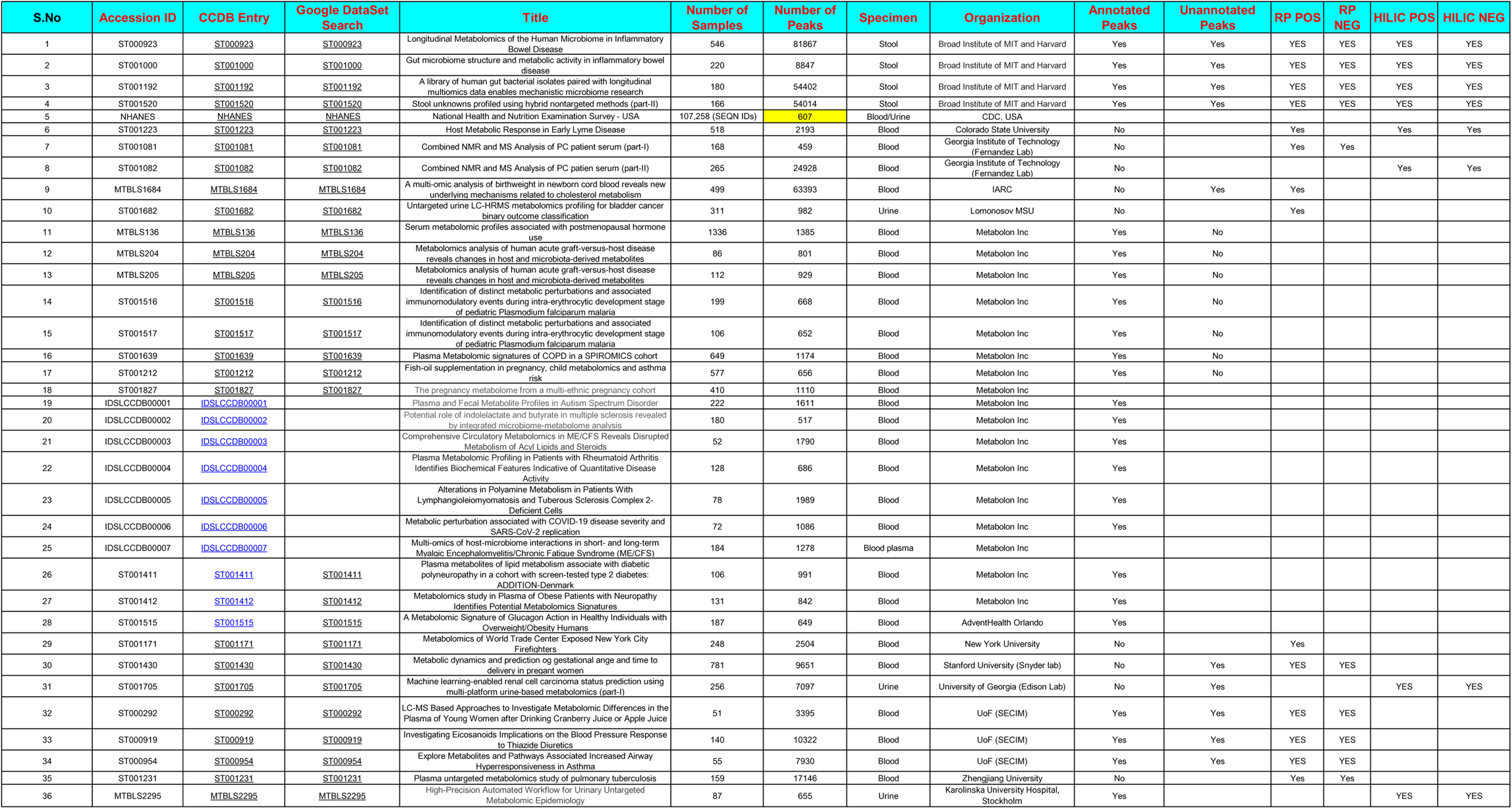
Description of the studies included IDSL.CCDB as of January 2022.

**Figure S1:**
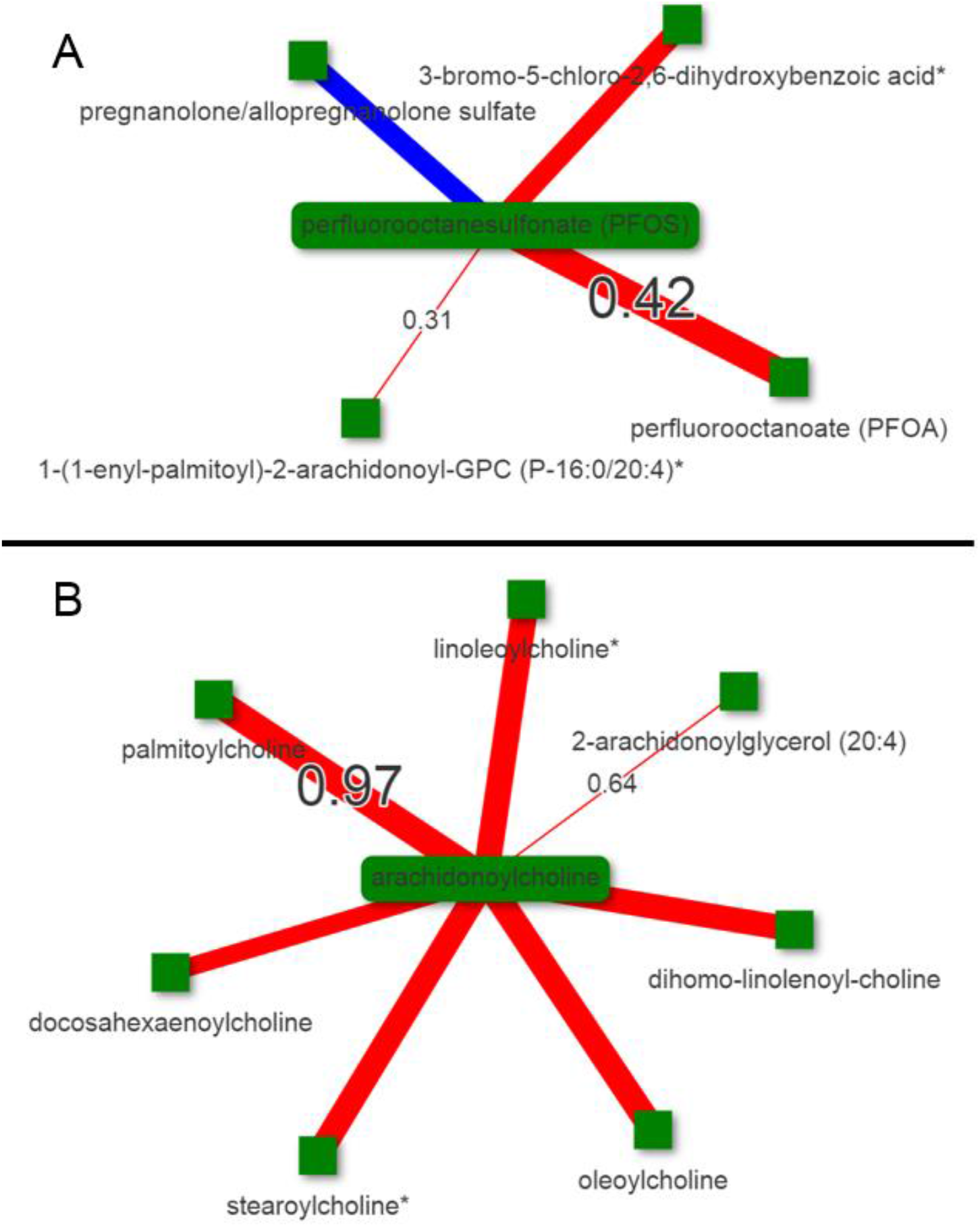
Exogenous compounds tend to have weaker correlations among similar chemicals in comparison to endogenous compounds. Only the minimum and maximum correlation values are highlighted on the edges for clarity. The study was https://chemcor.idsl.site/originaldata/metabolon/#?studyid=IDSLCCDB00007

**Figure S2.**
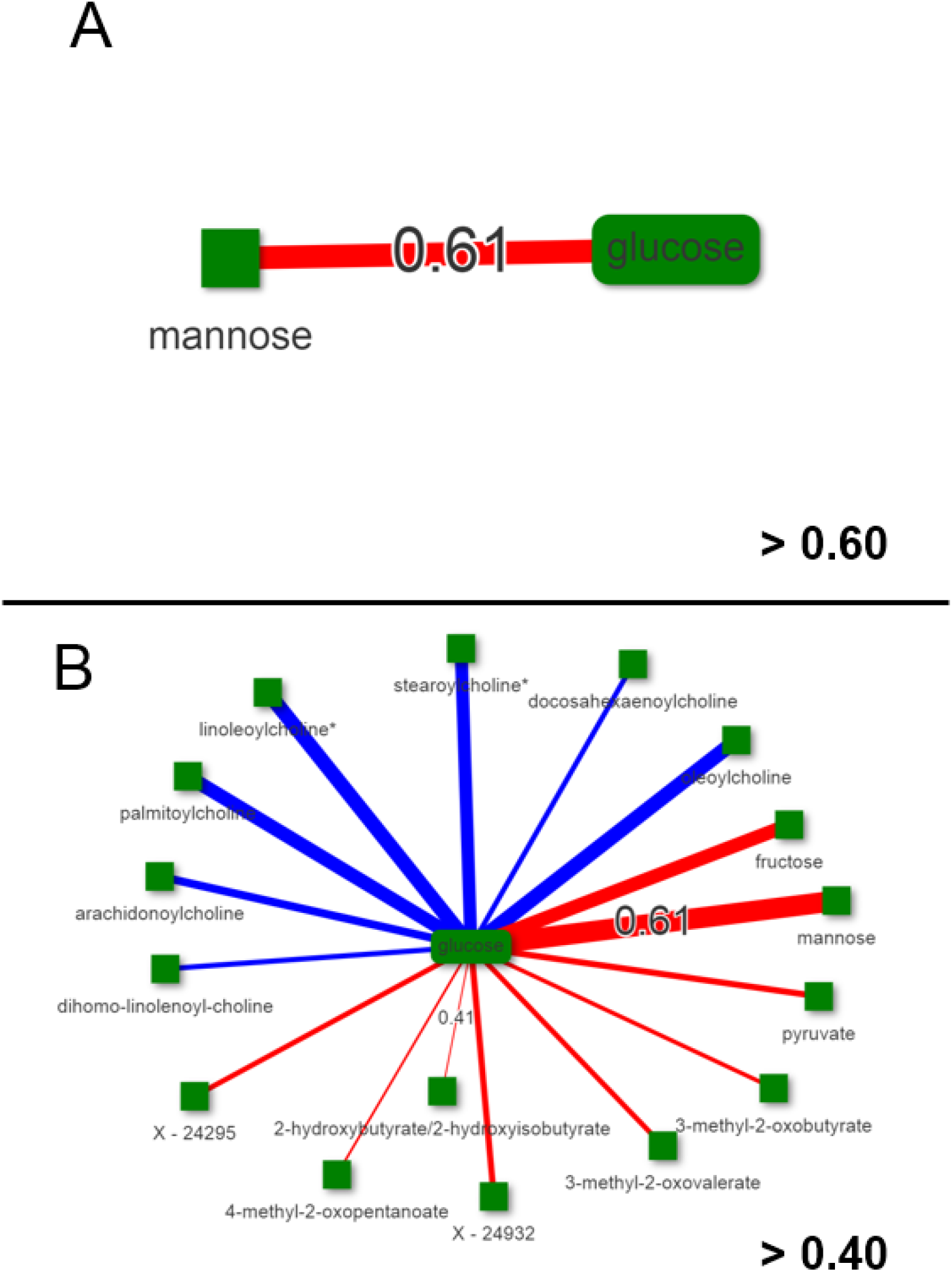
Only on a lower correlation threshold (0.40), metabolic relationships among glucose and acyl-choline metabolites were observed in the MTBLS136 study.

**Figure S3.**
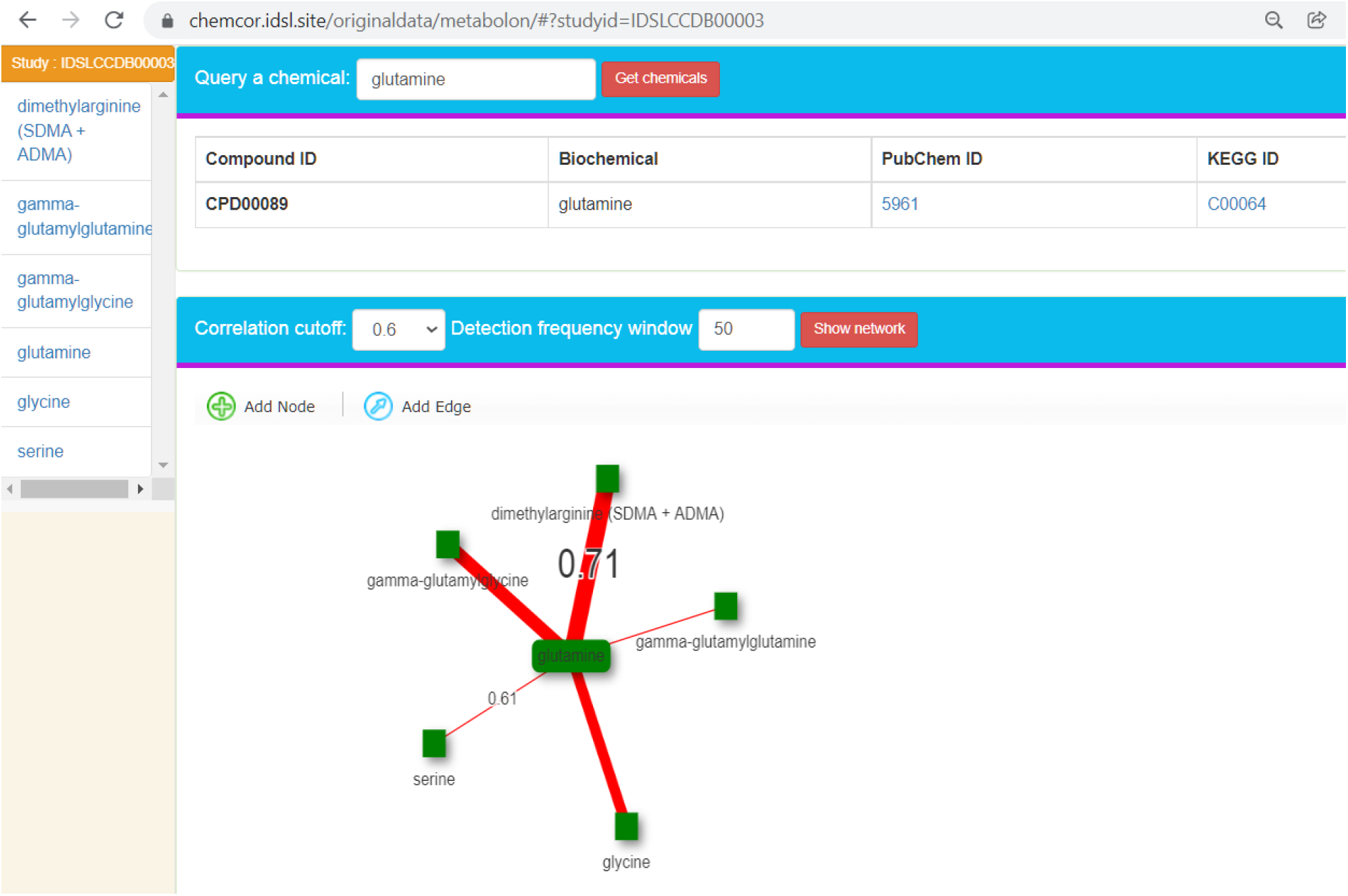
Web-interface for studies for which data were generated by the Metabolon Inc company.

**Figure S4.**
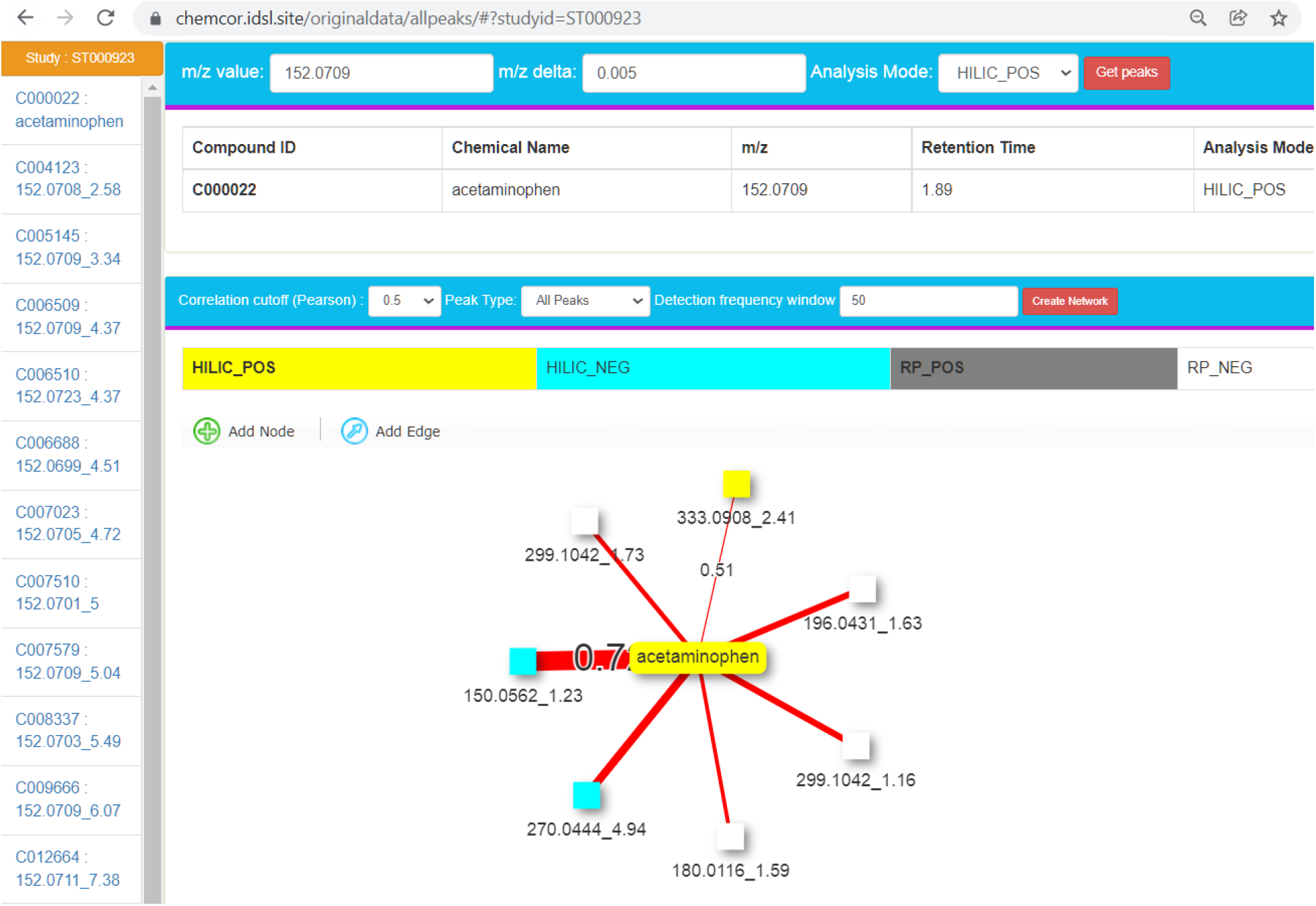
Web-interface for studies for which data were generated by an untargeted metabolomics assay using high-resolution mass spectrometry.

**Figure S5.**
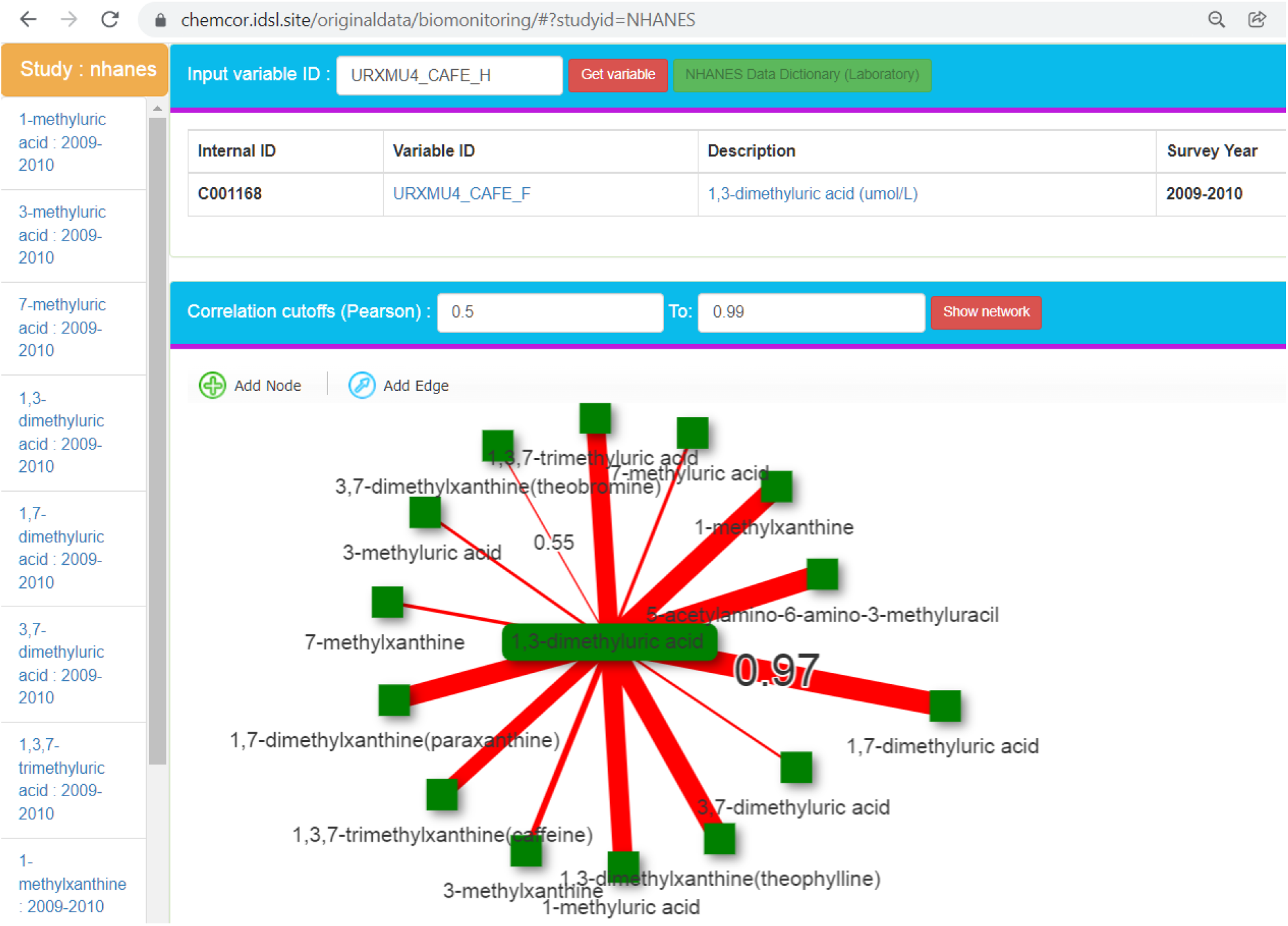
Web-interface for studies for which data were generated by the targeted assays that were used for the NHANES biomonitoring data.

**Figure S6:**
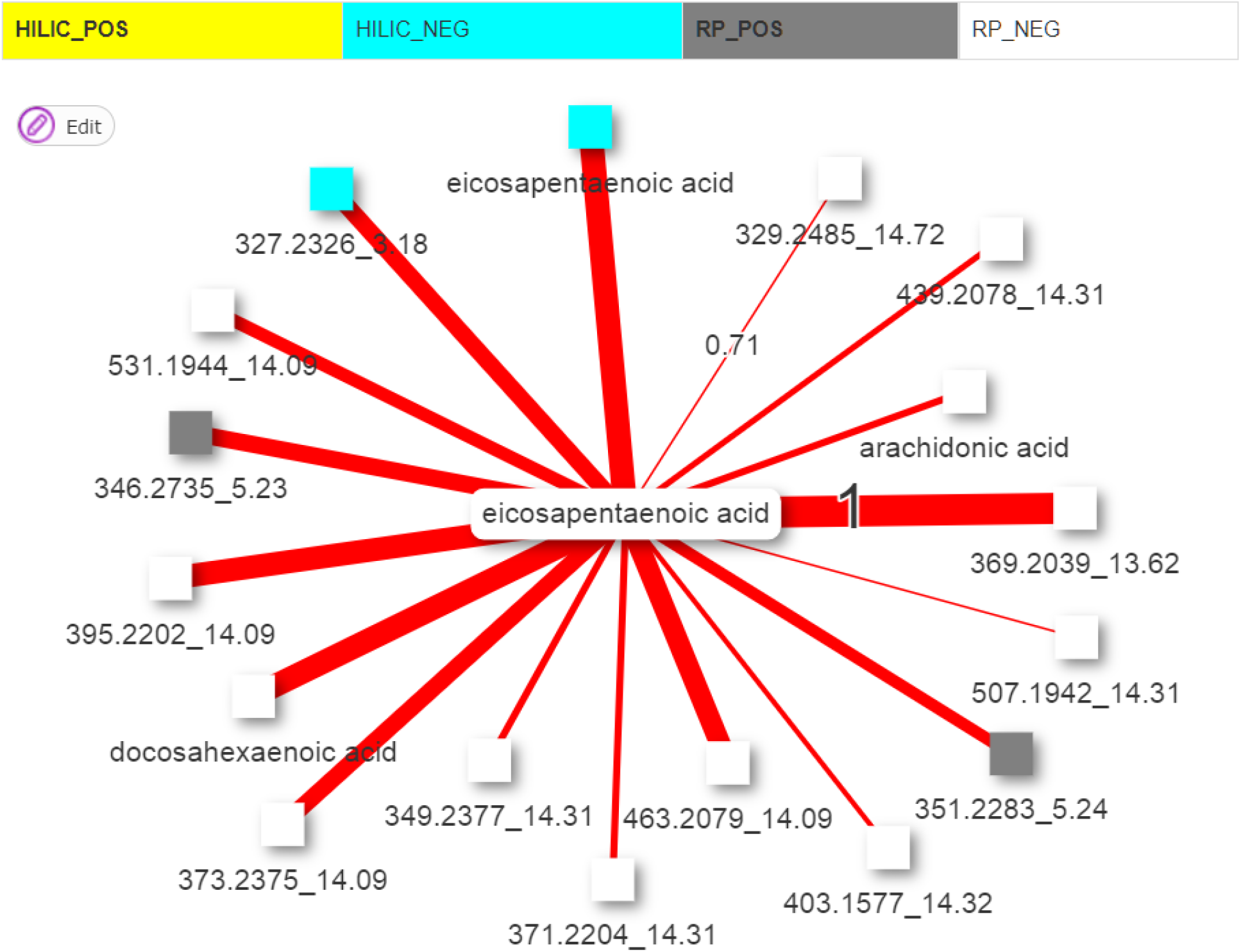
Compounds in two analysis modes. EPA was observed in both RP (−) and HILIC (−) modes in the study ST001000 so it showed a strong inter-chemical correlation. (https://chemcor.idsl.site/originaldata/allpeaks/#?studyid=ST001000)

**Figure S7:**
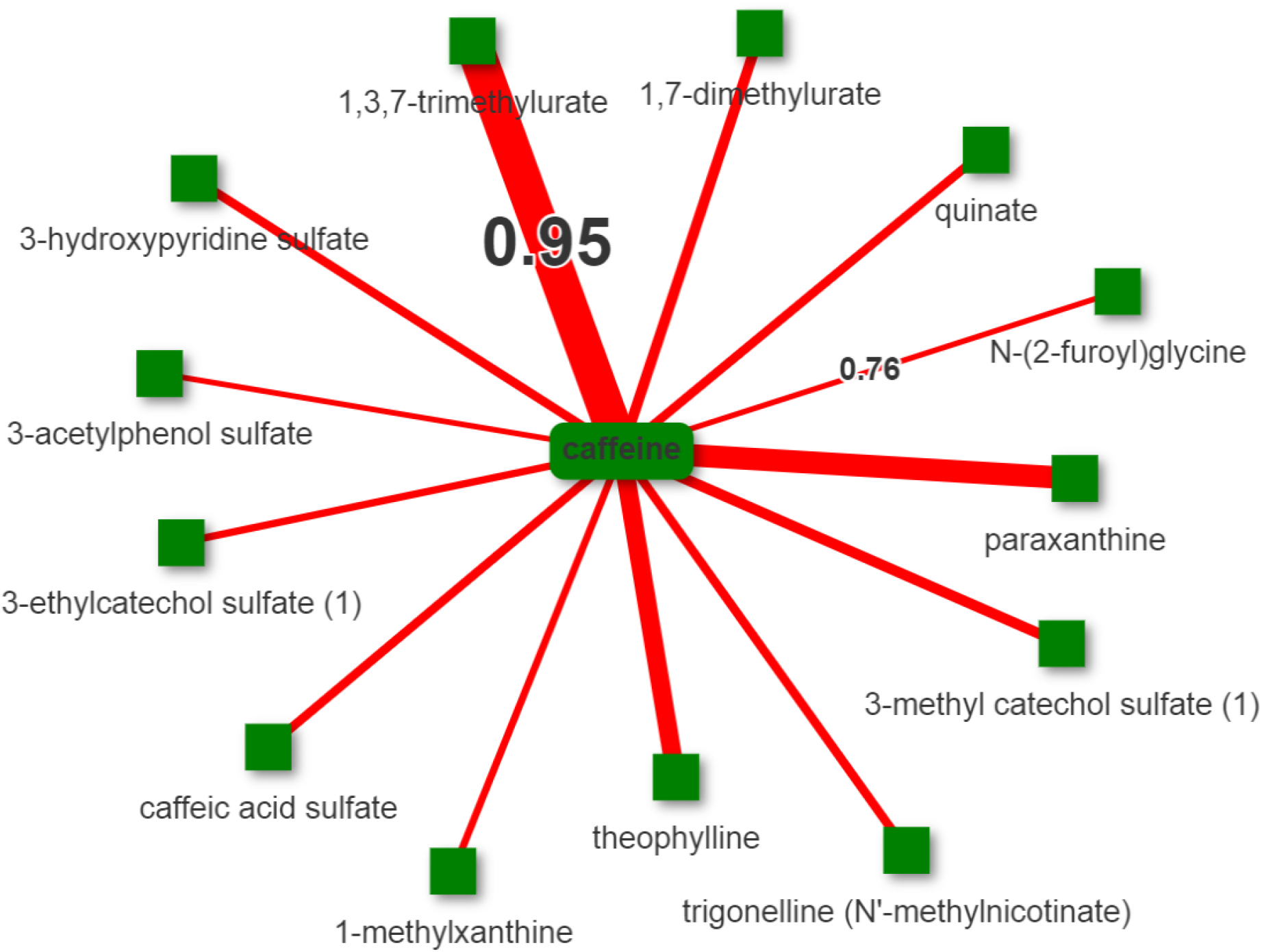
Caffeine and its correlating metabolites in the IDSLCCDB00007 study. Correlation threshold was 0.75. (https://chemcor.idsl.site/originaldata/metabolon/#?studyid=IDSLCCDB00007)

**Figure S8:**
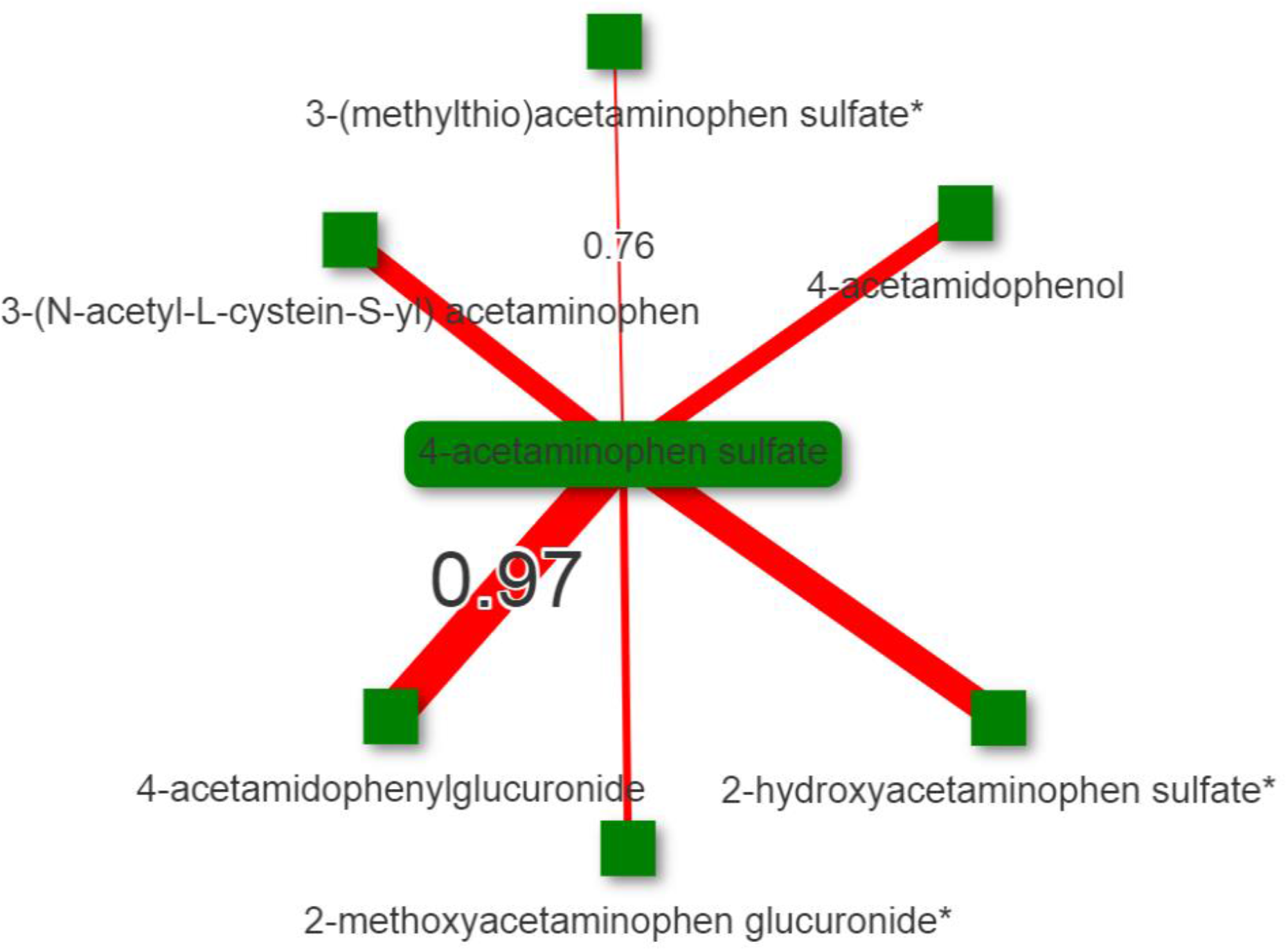
Acetaminophen metabolites showed strong inter-chemical correlations among them in the ST001639 study. https://chemcor.idsl.site/originaldata/metabolon/#?studyid=ST001639

